# Loss of α2,3-linked sialoside in the receptor-binding site of a H5N1 influenza hemagglutinin identified in a human patient

**DOI:** 10.64898/2026.01.19.700419

**Authors:** John H. Ni, Saeid Malek Zadeh, Alison M. Berezuk, Ryan Lynam, Peter Axerio-Cilies, Xing Zhu, Katharine S. Tuttle, Gethin Rh. Owen, Maria Tokuyama, Sriram Subramaniam

## Abstract

In November 2024, an adolescent female in British Columbia was hospitalized presenting with severe symptoms including respiratory failure due to infection with a novel H5N1 subtype influenza strain (BC24). Using cryogenic electron microscopy (cryo-EM), we show here that the N169 α2,3-linked auto-glycan that is found in the sialic acid binding site of previously studied H5 hemagglutinin (HA) proteins is absent in purified BC24 HA protein, suggesting greatly reduced affinity for α2,3-linked sialosides. Glycan microarray analysis shows that the BC24 HA protein displays reduced or no binding not just to most α2,3-linked sialosides, but also to α2,6-linked sialosides. Full-length BC24 HA expressed in A549 lung alveolar carcinoma cells drives membrane fusion, albeit at significantly lower levels than previous H5 HA proteins, and post-infection sera from the patient display strong binding to BC24 HA and HA proteins from other influenza subtypes. The high virulence of the BC24 strain despite weak receptor binding reveals further complexity in the factors that result in severe disease caused by avian influenza.

## INTRODUCTION

Outbreaks of H5N1 influenza in cattle and birds are a source of concern for zoonotic transmission to humans with a nearly 50% fatality rate in confirmed cases^1,2^. In November 2024, an adolescent female in British Columbia was hospitalized presenting with severe symptoms including respiratory failure and positivity for infection with novel H5N1 influenza in clade 2.3.4.4b (A/British Columbia/PHL-2032/2024)^3^, abbreviated as BC24 in this work. In addition to the well-studied E627K mutation in polymerase basic 2, the most frequent mutations observed in H5N1 sequences from this patient were E190D and Q226H (by H3 sequence numbering) in hemagglutinin (HA)^3^. HA is the surface glycoprotein necessary for influenza A infectivity, responsible for both binding host cell receptors and mediating fusion of the viral and host endosomal membranes following endocytosis of the infecting virion^4,5^.

The host receptors for HA are typically sialosides: those with α2,3-linked sialic acids (*N*-acetylneuraminic acid (Neu5Ac) in this study) are often considered avian, and those with α2,6 linkages are often considered human^4^. Both are present, however, in human airways, with the α2,6-linked sialosides tending to be enriched in the human upper respiratory epithelia, and α2,3-linked sialosides deeper in the respiratory system and in children^4^. The E190D mutation, in concert with G225D, was previously reported to be involved in H1 HA receptor switching to α2,6-linked sialoside binding and human infectivity^6,7^. The E190D mutation in isolation, however, does not appear to promote sialoside affinity in either H1 or H5 contexts^6–8^. Leucine at the 226 position in H5 HA has demonstrated α2,6-linked sialoside binding^5,8^; however, this effect was not found with the histidine at this position^5^.

To explore the role of these HA mutations observed in BC24 on structure, we carried out cryogenic electron microscopy (cryo-EM) of BC24 HA as well as the H5 HA proteins present in the commonly studied clade 1 A/Vietnam/1203/2004 (VN04) and the mild-human-disease-causing clade 2.3.4.4b A/Michigan/90/2024 (MI24). The latter strain is more recent and closely related to the sequences observed in the infected BC24 patient. We specifically assessed the BC24 HA carrying the sequence deposited in GenBank XTJ61994.1 with the two most commonly observed mutations, E190D and Q226H. To better understand the glycan-binding properties of these HA proteins, we assessed the binding of BC24 HA to a wide range of glycans in a microarray format. We also used enzyme-linked immunosorbent assays (ELISA) and fusion assays in A549 human epithelial carcinoma cells to determine the receptor binding effects of the individual E190D and Q226H mutations, and to evaluate the functional effects of the two mutations on HA-mediated membrane fusion under conditions that mimic the environment during infection.

## RESULTS

### Density for the N169 α2,3-linked auto-glycan is absent in BC24 HA

To determine whether the HA mutations in BC24 result in altered HA conformation, we determined the pre-fusion state cryo-EM structures of trimeric HA ectodomains for BC24, VN04, and MI24 expressed and purified under identical conditions (processing workflow in Supplementary Fig. 1-3). Globally, all HA proteins display similar tertiary and quaternary structures (Fig. 1a-c) with well-resolved fusion peptides sequestered in the helix C bundle (Supplementary Fig. 4). The structures demonstrate that the E190D and Q226H mutations do not impart substantial structural changes either close to (Fig. 1d-f), or distal to (Fig. 1a-c) the main residues mutated in BC24 HA.

**Figure 1.**
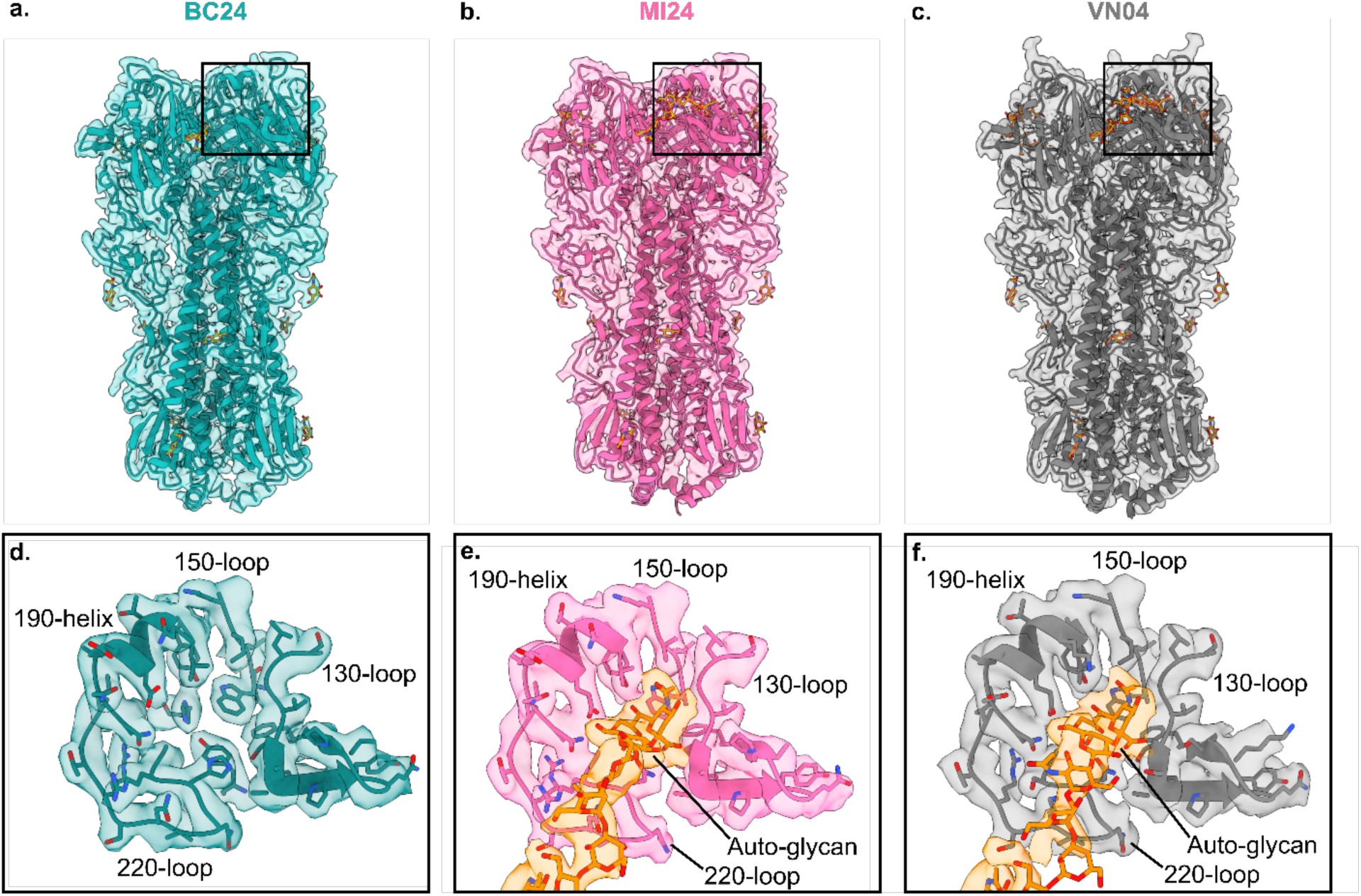
The BC24 HA receptor-binding site lacks auto-glycan density. From HA ectodomain constructs expressed in Expi293F cells, cryo-EM structures (map and model) are presented for **a.** BC24, **b.** MI24, and **c.** VN04 HA proteins. Glycosylation in all models is shown in orange. In each case, the black square indicates the region shown in panels **d-f**. To demonstrate loss of the α2,3-linked sialoside auto-glycan binding in the BC24 HA, maps and models of the receptor-binding sites are presented for **d.** BC24, **e.** MI24, and **f.** VN04 H5 HA proteins, with the auto-glycan in orange in each case.

The most striking finding from comparison of the different H5 HA structures is that clear cryo-EM density for the N169 auto-glycan is observed in the receptor-binding site for both the VN04 and MI24 HA proteins but not in BC24 HA (Fig. 1d-f). We modelled this full glycan as GlcNAc-GlcNAc-Man-Man-GlcNAc-Gal-Sia as previously reported by Morano et al. for a H5 HA with identical sequence to MI24 HA expressed in Expi293F cells (Fig. 1e,f)^9^. This glycan is likely branched^10^, but the branch not bound by the receptor-binding site remains too flexible to resolve. In the normal course of events, these auto-glycans are displaced by sialosides present on the host cell membrane for infection. Asparagine at the 169 position is highly conserved across H5 HA proteins^9^, including in the BC24 and VN04 HA proteins. The density maps for VN04 and MI24 HA permit confident modelling of the first two sialic acid and galactose monomers (Sia-1 and Gal-2), showing the terminal glycosidic linkage being in the α2,3 configuration (Fig. 1e,f), distinct from the geometry of the α2,6-linkage^8^. The finding that the BC24 HA receptor-binding site is not normally occupied by the auto-glycan suggests that this HA is likely to have reduced affinity for binding α2,3-linked sialosides on the host cell to initiate viral entry.

### Comparison of glycosylation patterns in BC24 and other H5 HA proteins

HA is known to be extensively glycosylated at multiple sites^11^. We compared the densities for *N*-linked glycans between the different HA proteins to assess whether there were any differences in glycosylation patterns, and to determine whether the absence of auto-glycan in the receptor-binding site is due to lack of glycosylation at N169 in the BC24 HA protein. The cryo-EM reconstructions show density in the three HA proteins extending from *N*-linked glycosylation sites at positions N21, N33, N158 (VN04 only), N169, N289, and N483 (Fig. 2a-f). We were able to model the first portion at N33, N169, N289, and N483 with sufficient resolution (Fig. 2b,d-f). For N169, density beyond the initial portion is resolved only with HA from VN04 and MI24, where these glycans are bound at the receptor binding site (Fig. 2d). The lack of density in the BC24 receptor-binding site (Fig. 1d) is therefore not due to lack of glycosylation at N169, but because of the reduced affinity for this glycan to the binding site. Density connecting glycans to two other residues (N21 and N158, the latter only in VN04) is also visible, but the glycans themselves are likely disordered and could not be modeled reliably (Fig. 2a,c). The presence of the glycan at N158 had previously been reported to moderately mask the receptor-binding site, affecting receptor tropism, and the clade 2.3.4.4b HA sequences lack this glycosylation site, potentially contributing to human infectivity^12^.

**Figure 2.**
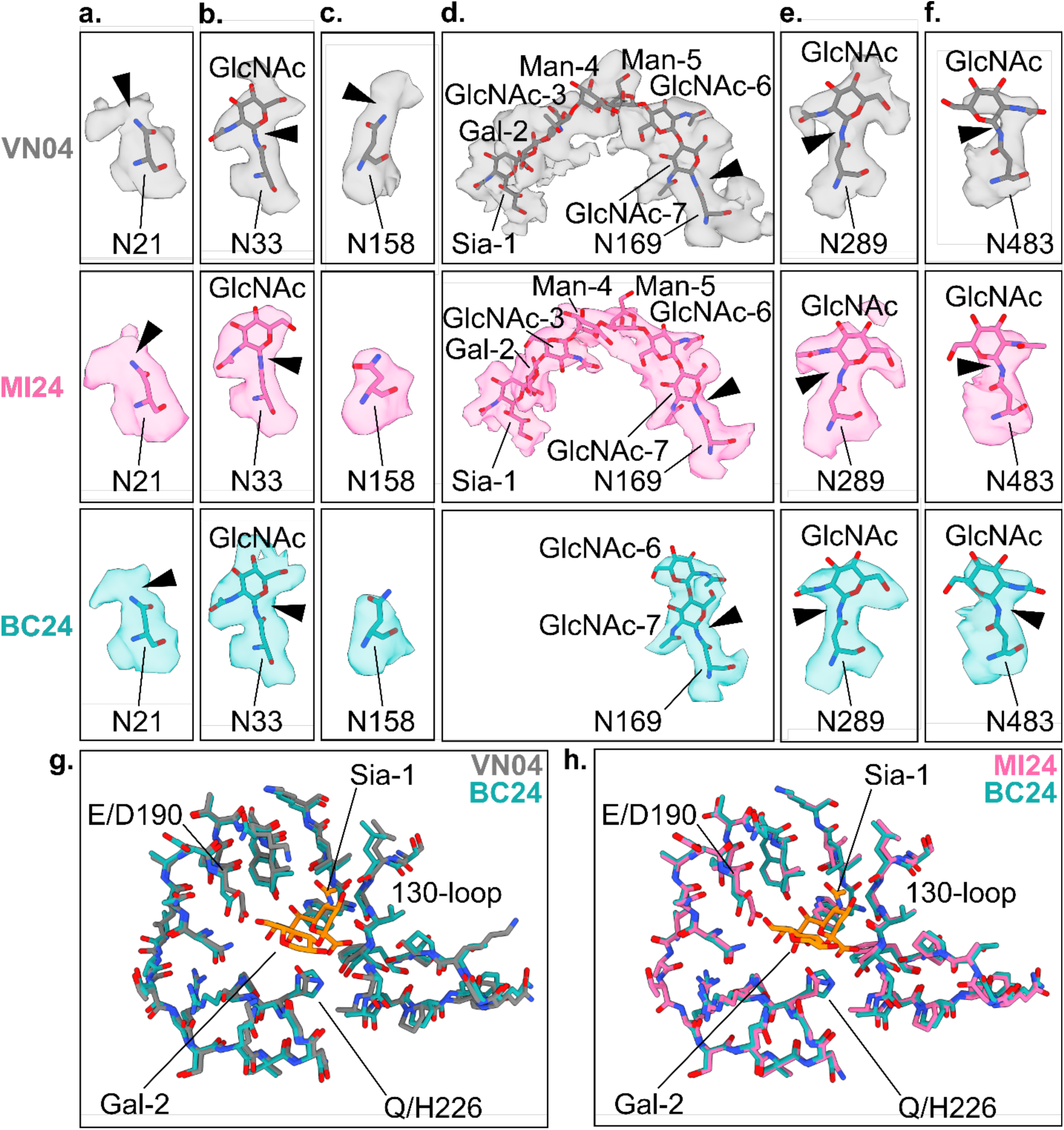
Glycosylation present in H5 HA proteins. Zoomed-in views of cryo-EM structures (map and model) are presented at the following *N*-glycosylation sites: **a.** N21, **b.** N33, **c.** N158, **d.** N169, **e.** N289, and **f.** N483. Top, VN04 HA; middle, MI24 HA; and bottom, BC24 HA. Arrowheads indicate junctions between glycan and protein density. Sia, sialic acid (Neu5Ac); Gal, galactose; GlcNAc, *N*-acetyl glucosamine; and Man, mannose. Locally aligned cryo-EM models of receptor-binding sites are presented for **g.** VN04 and BC24 HA proteins and **h.** MI24 and BC24 HA proteins. Teal, BC24; grey, VN04; and pink, MI24. The auto-glycan (in orange) from VN04 and MI24 structures is shown with only the first two residues for clarity.

### Structural explanation for the lack of bound α2,3-linked auto-glycan in BC24 HA

Alignment of the receptor-binding site residues also allows us to compare the bound and unbound structures across these differing H5 HA proteins. The receptor-binding site structure is relatively unchanged in the three H5 HA proteins (Fig. 2g,h). In the VN04 and MI24 HA structures, the E190 carboxylate is positioned for hydrogen bonding 2.9 and 2.8 Å, respectively, from the Sia-1 C9 hydroxyl. Their Q226 amides are respectively 3.3 and 3.2 Å from the Gal-2 C4 hydroxyl, and the MI24 Q226 amide is also 3.2 Å from the glycosidic bond oxygen linking Gal-2 and Sia-1. In the absence of bound glycan, the BC24 D190 carboxylate, spaced from the backbone by one fewer methylene moieties, is positioned 4.9 Å from where the MI24-bound Sia-1 aligns. Similarly, H226 of BC24 is pivoted 1.1 Å away from the position of the MI24-bound Gal-2 C4 hydroxyl and 0.9 Å towards the C1 carboxylate, likely permitted by the lack of ligand in the pocket. Thus, key residues in HA that appear well-suited in VN04 and MI24 HA to interact favorably with the α2,3-sialylated N169 auto-glycan are also the ones mutated in BC24 HA suggesting a structural explanation for the observed lack of α2,3-linked glycan density in the sialic acid binding pocket.

### Glycan microarray analysis

With structural evidence that the BC24 HA may have significantly reduced α2,3-linked sialoside binding, we sought to understand the glycan binding profiles of VN04, MI24, and BC24 HA proteins through microarray analysis featuring a range of *N*-linked glycans. The VN04 and MI24 HA proteins demonstrated binding exclusively to α2,3-linked sialosides (Fig. 3a,b). The MI24 HA bound to a greater breadth of α2,3-linked sialosides and tended to bind more strongly than VN04 HA (Fig. 3a,b). The MI24 HA had a mild binding preference for sialyl Lewis^X^ in particular (Fig. 3b). A weak but noticeable signal was observed for both of the clade 2.3.4.4b MI24 and BC24 HA proteins to the core Man_3_GlcNAc_2_ glycan (Fig. 3b,c). Consistent with the cryo-EM structural findings (Fig. 1), the BC24 HA did not demonstrate binding to any of the α2,3-linked sialosides or any of the glycans represented on the microarray (Fig. 3c).

**Figure 3.**
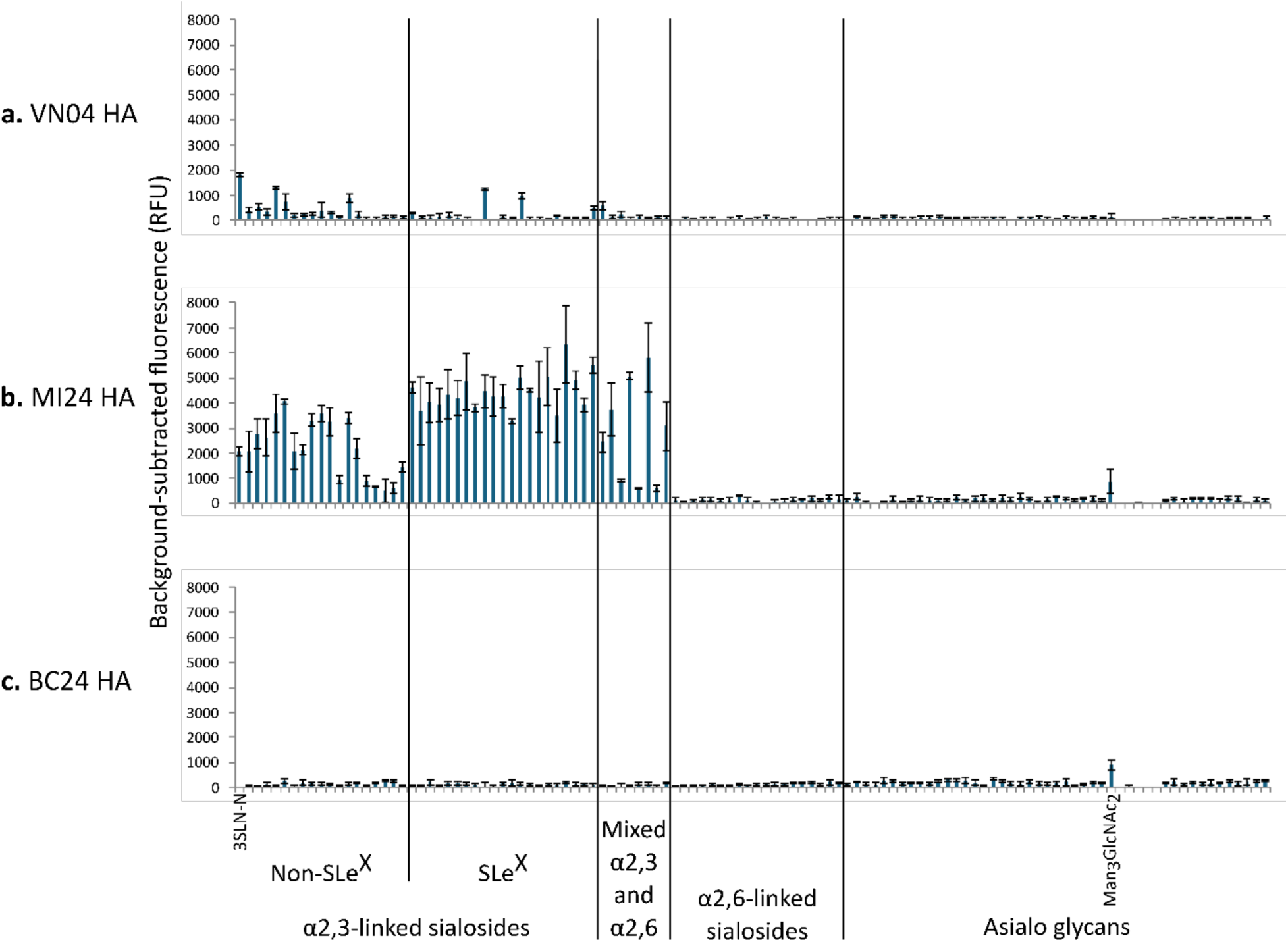
*N*-glycan microarray analysis of H5 HA proteins. HA expressed from GnTI-Expi293F cells were precomplexed for 1 h on ice with anti-6xHis primary antibody and Cy3-conjugated secondary antibody at a 10:1:1 ratio. This precomplex solution was then allowed to incubate on a blocked *N*-glycan microarray slide for 1 h at room temperature before scanning the slide at 532 nm. Data presented have signal from control spots without glycan subtracted. **a.** VN04 HA, **b.** MI24 HA, and **c.** BC24 HA. Bar height represents the mean, and error bars are standard deviation. SLe^X^, sialyl Lewis^X^.

The weaker binding of VN04 HA compared to MI24 HA across tested sialosides is consistent with earlier studies of sugar binding to HA proteins. Pulit-Penaloza et al. reported that the VN04 HA did not bind to most fucosylated glycans whereas the MI24 HA bound well to these glycans, which is reproduced in our results (Fig. 3a,b)^13^. Their findings show that the strongest binding of the VN04 and MI24 HA proteins is to sulfated glycans which are not represented on our microarray^13^. Furthermore, binding to 3SLN-N (leftmost bar of Fig. 3) was comparable between MI24 and VN04 HA proteins in this assay, so it is possible that VN04 HA may bind better to the *O*-linked glycans represented in previously reported microarray analyses^7,13^. We performed dynamic light scattering (DLS) analysis showing that all protein preparations used in the microarray analysis are comparable in stability (Supplementary Fig. 5), which indicates that the differences in binding profiles are intrinsic to the receptor selectivity of the HA proteins analyzed.

### Assessment of glycan binding by E190D and Q226H HA mutants using ELISA experiments

With the reported mixed population of receptor-binding site mutations in sequencing^3^, we generated single and double E190D and Q226H mutants on the MI24 HA background to investigate their individual roles in receptor binding and/or synergy between these mutations. Using ELISA measurements, we subsequently assessed binding of these HA mutants to typical α2,3-linked sialosides 3SLN-N (represented on the microarray as N1, leftmost bar in Fig. 3) and 3SLN_3_-N and typical α2,6-linked sialosides 6SLN-N and 6SLN_3_-N, noting that previous studies have identified that longer sugars improve HA binding affinity^8^. Consistent with the microarray findings, we observed concentration-dependent binding of both VN04 and MI24 HA to the α2,3-linked sialosides but not to the α2,6-linked sialosides (Fig. 4a,b). Binding to the longer glycan was stronger for both of these HA proteins (Fig. 4a,b). Neither the BC24 HA nor any of the related receptor-binding site mutants (with MI24 background for the rest of HA) showed significant binding to either sialoside variety within the tested range up to 50 μg/mL (Fig. 4c-f). We verified the robustness of these results with a biological replicate of the assay using separate preparations of each HA ectodomain (Supplementary Fig. 6). Taken together, both the E190D and Q226H mutations on MI24 HA background individually reproduce the BC24 HA behavior of strongly reduced binding to both α2,3-linked and α2,6-linked sialosides.

**Figure 4.**
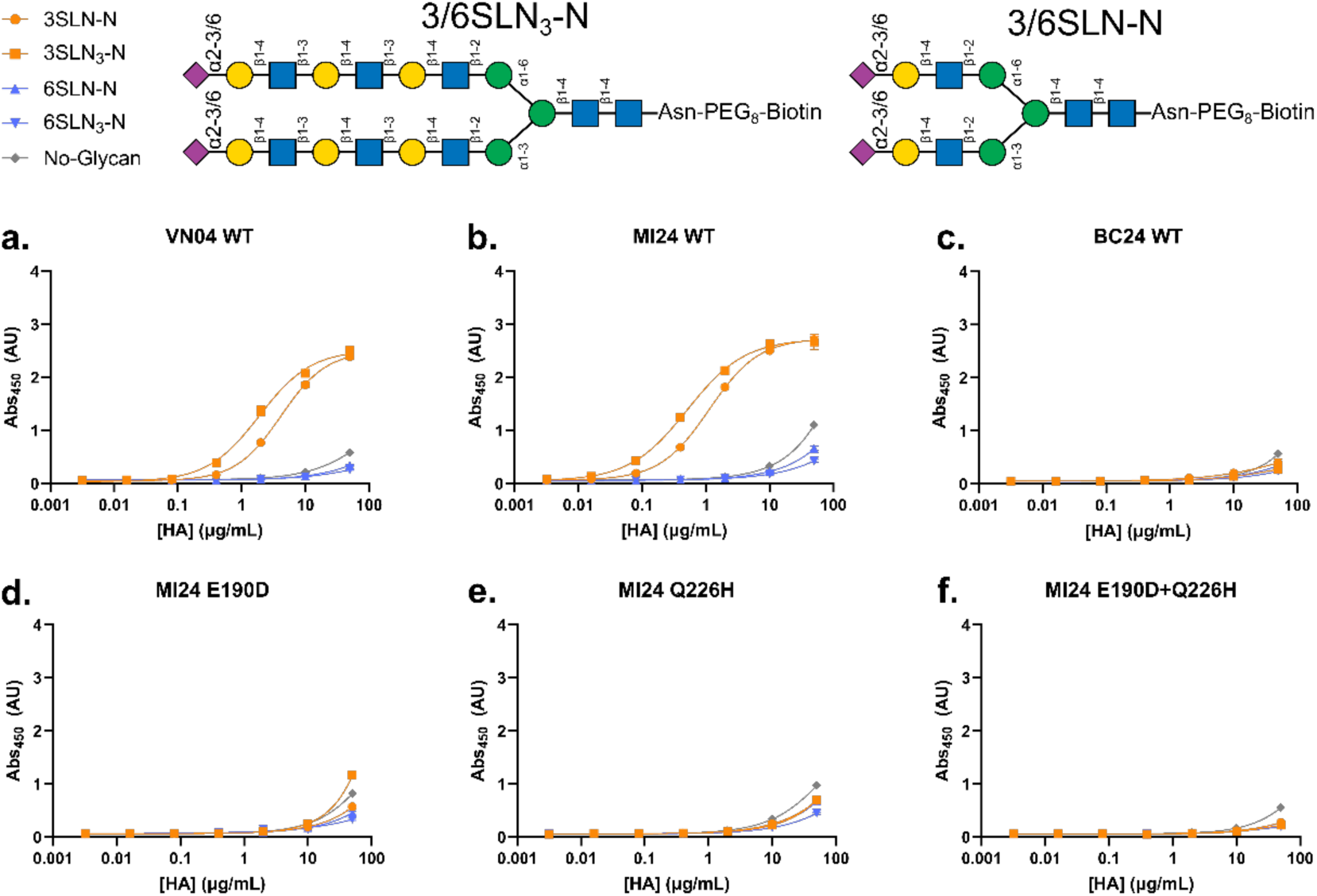
Individual BC24 HA receptor-binding mutations reduce binding to α2,3- and α2,6-linked sialosides. Representative ELISA curves probing binding at room temperature for 2 h to biantennary α2,3- and α2,6-linked sialosides (schematic at top) by the following HAs: **a.** VN04 wild-type, **b.** MI24 wild-type, **c.** BC24 wild-type, **d.** MI24 E190D mutant, **e.** MI24 Q226H mutant, and **f.** MI24 E190D+Q226H double mutant. Error bars are standard deviation (n=3).

### BC24 HA is competent for membrane fusion

Since BC24 HA permitted infection of the patient despite the apparently reduced affinity to the assayed glycans, we investigated HA-driven membrane fusion activity in A549 alveolar carcinoma cells transfected with full-length VN04, MI24, or BC24 HA proteins. The transfected cells display HA on the plasma membrane and can be activated to trigger cell-cell fusion by transiently lowering the extracellular pH to mimic the low pH environment of late endosomal membranes. Membrane fusion generates multinucleate syncytia and mixing of the cytoplasm of cell pairs permitting reconstitution of functional luciferase from cells separately transfected with fluorescently tagged NanoBiT subunits. The subsequent luminescence therefore indicates the amount of fusion events that occurred. We performed western blotting to verify that variations in luminescence were not due to differential HA expression (Supplementary Fig. 7).

In agreement with the published evidence for BC24 HA function in the case report^3^, fusion events are clearly observed in A549 cells transfected with all of the HA proteins, including BC24, by fluorescence microscopy (Fig. 5a). We then quantified the extent of fusion by probing the activity of reconstituted split NanoBiT luciferase (Fig. 5c). Statistically significant increases in luminescence relative to mock-transfected controls were observed in all HA-transfected cultures (Fig. 5c and Supplementary Table 1). The levels of fusion mediated by the BC24 HA were substantially lower than by VN04 and MI24 HA (Fig. 5c). Given the absence of glycan binding by BC24 HA in the structural and glycan array binding experiments, it is possible that fusion events with BC24 HA may be driven by avidity effects, compensating for the low biochemical affinity^14^.

**Figure 5.**
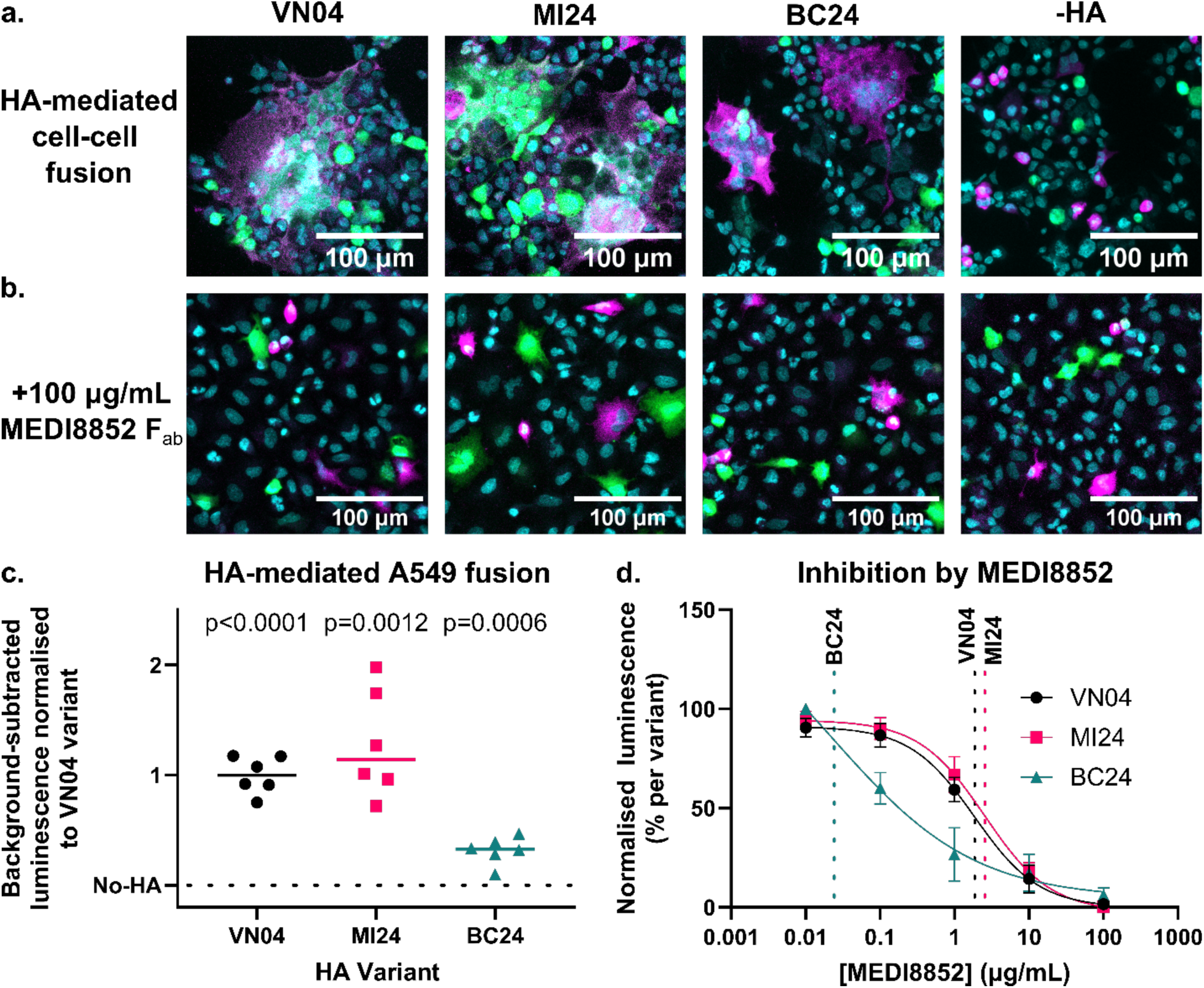
BC24 HA is fusion competent and neutralizable by MEDI8852. A549 human alveolar carcinoma cells were transfected separately with GFP-HiBiT, or both HA and mCherry-lgBiT. HA and mCherry-lgBiT cotransfected cells were then overlaid on GFP-HiBiT cells and acidified at pH 4.5 for 3 min. Following recovery, luciferase activity was assessed as a proxy for fusion. Cells were imaged following fixation in 4% paraformaldehyde and counterstaining in DAPI. Green, GFP; magenta, mCherry; and cyan, DAPI. **a.** Representative micrographs showing syncytia uniquely in HA-transfected conditions. **b.** Inhibition of fusion by 1 h incubation with MEDI8852 F_ab_ prior to acidification. **c.** Summary of fusion efficiency by each HA. Data presented have background signal from mock transfected controls subtracted and are subsequently normalized to the signal from the VN04 HA within each set of biological replicates. Horizontal lines represent the median. Significance values are against mock transfected control. **d.** Dose-inhibition curves of MEDI8852 F_ab_ for each HA with y-axis normalised to be the range of observed signal for each HA. Vertical dotted lines represent the calculated IC_50_. Error bars are standard error of the mean (n=6).

MEDI8852 is a broadly neutralizing antibody which recognizes a site on HA that is conserved in most HA subtypes^15^. To test whether BC24 was susceptible to neutralization by MEDI8852, we repeated fusion assays with a dilution series of MEDI8852 F_ab_ prior to HA activation to estimate the extent to which BC24 HA activity can be neutralized by this antibody (Fig. 5b,d). IC_50_ values computed based on NanoBiT luciferase assays demonstrate that BC24 is potently neutralized by MEDI8852 at least as effectively as VN04 and MI24 (estimated IC_50_ of 0.502 nM for BC24 HA versus 38.6 nM and 52.9 nM for VN04 and MI24 HA, respectively). Biolayer-interferometry (BLI) experiments with MEDI8852 F_ab_ validate specific binding in the nanomolar range (Supplementary Table 2 and Supplementary Fig. 8) with all three HA proteins.

### Humoral immune response against BC24 H5 HA

Based on our findings that the severity of disease for the BC24 case is not coupled to strong HA binding to α2,3- or α2,6-linked sialosides and that BC24 H5 HA caused fewer fusion events, we sought to determine the BC24 patient’s antibody titer against BC24 H5 HA. We tested serum samples obtained from the BC24 patient at four independent time points within 1 to 4 days following negative serum Flu A RT-PCR test results^3^ and from four age-matched healthy controls (Supplementary Table 3). Titration of serum antibodies against BC24 and MI24 HA proteins showed that healthy controls have modest reactivity against both of these clade 2.3.4.4b H5 HA proteins (Fig. 6a). Since these individuals have no known prior history of exposure to H5N1 influenza, these are likely cross-reactive antibodies generated through vaccination or in response to circulating strains of influenza. In contrast, serum from the BC24 patient demonstrates strong reactivity against both the BC24 and MI24 HA proteins (Fig. 6a), indicating that the patient mounted a robust antibody response to the BC24 H5 HA protein.

**Figure 6.**
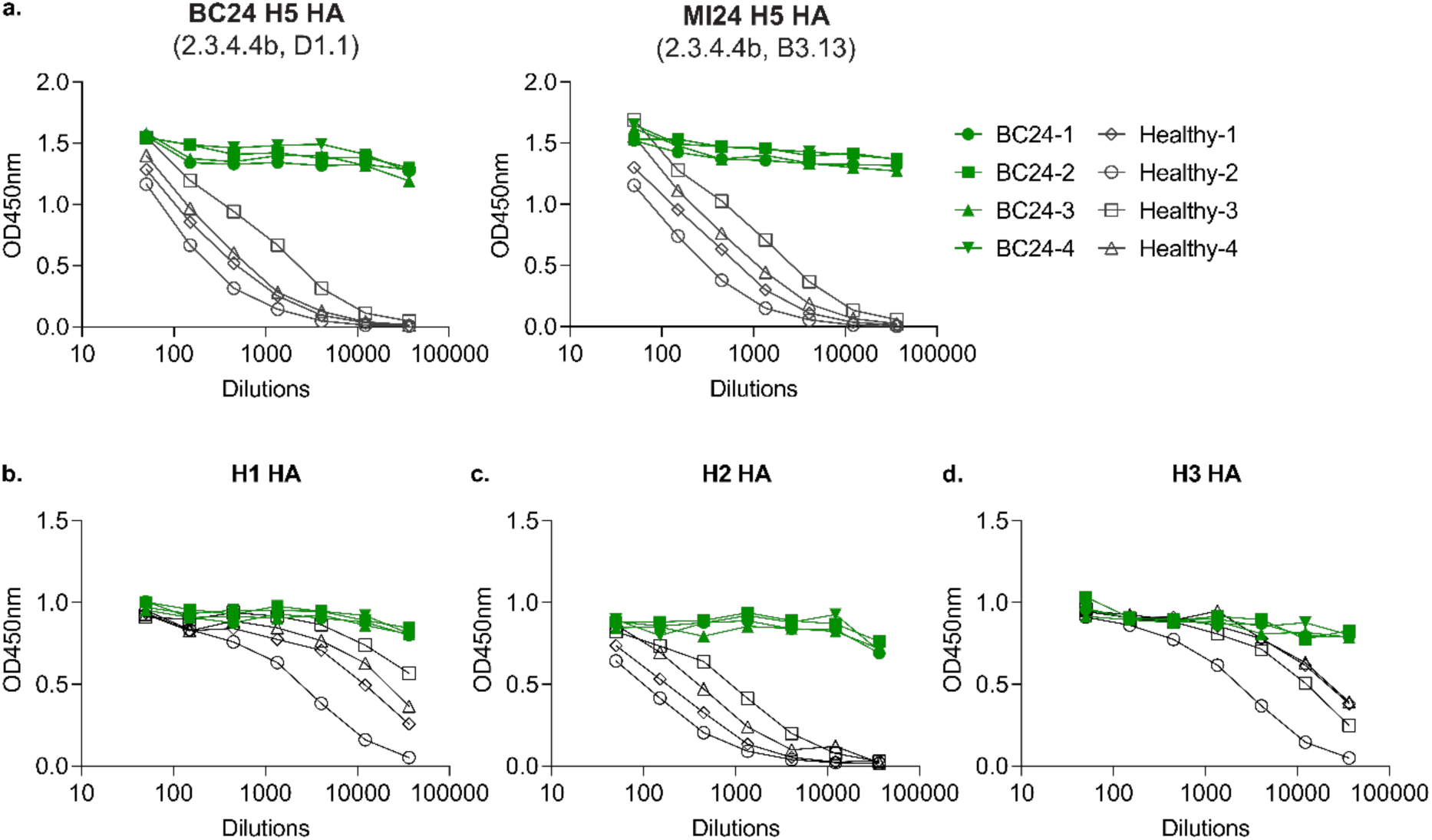
Patient serum reactivity against BC24 and MI24 HA measured by ELISA. **a.** BC24 (left) and MI24 (right) HA, **b.** H1 HA, **c.** H2 HA, and **d.** H3 HA proteins were incubated with serially diluted serum samples (green, sera from the BC24 patient collected at four different time points immediately following negative serum Flu A RT-PCR test result; grey, sera from unremarkable age-matched healthy donors). The data represent background subtracted OD450nm measurements.

To determine whether this reactivity was specific to H5 HA, we repeated the ELISA experiment against HA proteins from A/California/04/2009 (H1N1), A/Singapore/1/1957 (H2N2), and A/Perth/16/2009 (H3N2). As expected, the healthy controls showed a robust response against H1 and H3 HA proteins, which are part of circulating strains of influenza A and incorporated in seasonal vaccines, and mild reactivity to H2 HA (Fig. 6b-d). Notably, sera from the BC24 patient reacted strongly against all three HA proteins, including H2 HA even at the highest dilution (Fig. 6b-d) despite the negative nasopharyngeal swab obtained for A(H1) and A(H3) viral infection at the time of admission.

## DISCUSSION

### Implications for host receptor binding by BC24 HA

Our structural, biochemical, and cellular analyses collectively demonstrate diminished affinity of BC24 HA to α2,3-linked sialosides on host cells. The structures of several H5 HA proteins reveal a lack of BC24 HA binding to the auto-glycan (Fig. 1d). In contrast, the MI24 HA which differs from BC24 HA by only nine residues in the ectodomain per protomer, and by only E190D and Q226H amongst receptor-binding site residues, is competent to bind the auto-glycan, showing continuous glycan density even in the absence of stabilization by antibodies^9^. Given the similarity in receptor-binding site conformations between the BC24, VN04, and MI24 HA proteins (Fig. 2a,b), the reduced auto-glycan affinity likely arises from lack of specific interactions with the altered 190 and 226 positions, where E190 and Q226 appear to be favorable to binding in other strains.

The glycan microarray and ELISA experiments further demonstrate that the BC24 HA does not detectably bind the assayed α2,6-linked sialosides (Fig. 3c and 4c). Indeed, the mutagenesis experiments with ELISA-based readout suggest that each of the E190D and Q226H mutations yield reduced affinity for sialosides in H5 HA (Fig. 4d-f). Our cell-based assay showing reduced fusion in the BC24 HA-transfected cultures when compared to VN04 and MI24 HA further supports the interpretation that affinity for the most common glycans is reduced (Fig. 5c). The A549 alveolar cells used in this assay have been previously demonstrated to express both α2,3-and α2,6-linked sialosides on their surface^16^. Our findings therefore do not support the hypothesis^3^ that the mutations at residues 190 and 226 may have led to a switch to in HA receptor specificity from α2,3-linked to α2,6-linked sialosides.

Interestingly, the HA protein (MI24) which demonstrated the strongest binding to sialosides in the microarray analysis (Fig. 3b) and mediated the highest levels of A549 cell fusion (Fig. 5c) was identified from the influenza case with the mildest symptoms, limited to conjunctivitis^17^. By contrast, the VN04 and BC24 cases were respectively fatal and severe^3,18^, while demonstrating weaker receptor binding (Fig. 3a,c), although there are biologically relevant glycans not explored in this study^7,13^. We propose that the reduction in binding affinity, with retention of some level of fusion competence (Fig. 5c), may have allowed BC24 influenza to avoid receptors present in the upper respiratory tract. The BC24 virus may have thus penetrated more deeply into the respiratory system, resulting in more severe disease. Supporting this hypothesis, the patient had positive Flu A RT-PCR results for a longer time for samples from tracheal aspirate compared to throat swabs, and higher viral loads in the lower respiratory tract as compared to the upper respiratory tract^3^. Oner et al. had also found in the H5N1 outbreaks in the 2000s that presentation with rhinorrhea correlated with milder disease, similarly speculating on a protective effect by not permitting infection of the lower respiratory system^19^. Although the specific route of infection is unknown, given that the first symptoms that arose in the BC24 case were conjunctivitis and fever, it is possible that the MI24 and BC24 influenzae may have used the eye as a site to initiate infection^3,17^; however, the MI24 influenza may have been more confined to the eye by binding better to the α2,3-linked sialosides expressed there^20^.

The microarray used in this study does not include sulfated glycans which are displayed on respiratory cells^4^. These glycans have previously been shown to bind to H5 HA proteins^13^. Interestingly, H1 HA proteins were found to bind sulfated α2,3-linked sialosides with both aspartate or glutamate at position 190 while the non-sulfated α2,3-linked sialoside binding was hindered by introducing the E190D mutation^7^. It is possible that BC24 may use other glycans than those tested in our microarray experiments for host cell recognition.

The BC24 case did not result in any subsequent human-to-human transmission, which is consistent with the prevailing understanding that α2,6-linked sialoside binding promotes human spread^21,22^. However, this BC24 strain presents severe disease at least at the individual health level despite lacking well-understood human disease markers like increased α2,6-linked sialoside affinity in the HA protein. Ultimately, the adaptation of viruses to human infectivity requires further study to unveil such nuances. Fusion activity-based screening may provide an additional important approach to better understand zoonotic potential in surveillance efforts.

### Potential non-viral explanations for the severity of the infection

Pending further investigation of the genes sequenced from the BC24 strains, explanations for the BC24 case include host factors that are independent of the nature of HA. Patient age may have played a role, noting that teen-aged children have historically been particularly susceptible to severe disease attributed to H5N1 influenza due to lack of pre-existing immunity and different receptor expression profiles^4,19,23^. Factors such as asthma may also have played a role in the severity of the infection^3,24–26^. Although a specific understanding of the underlying causes differs between studies, with environmental factors often being substantial covariates, hyperresponsive immune systems are a consideration in the study of infections in patients with asthma^3,24–27^.

### BC24 influenza is susceptible to existing therapeutics

Although the BC24 influenza was capable of causing severe disease, both of the recent BC24 and MI24 HA proteins remain as readily neutralized by MEDI8852 as the VN04 HA (Fig. 5d). Most clade 2.3.4.4b H5N1 continue to be susceptible to commonly licensed anti-influenza drugs^28^, and the neuraminidase gene sequenced from this patient does not carry the H275Y (N1 numbering) mutation circulating in British Columbia around the time^29^, nor any other known resistance mutations^28,30^. Patient samples demonstrated susceptibility to oseltamivir^3^. Concerns of near-term health emergencies from zoonotic transmissions of related flu strains are largely assuaged because of the lack of resistance of these viral strains to traditional antiviral methods, and because the HA itself retains the conserved stem region that is efficaciously targeted by broadly-neutralizing antibodies (Fig. 5b,d; Fig. 6; and Supplementary Fig. 8).

### Humoral immune response against clade 2.3.4.4b H5 HA

The serum ELISA experiments demonstrated that all four healthy pediatric samples had detectable antibodies against clade 2.3.4.4b BC24 and MI24 HA proteins (Fig. 6a), likely due to cross-reactive antibodies from pre-existing immunity against vaccine or circulating strains of influenza A. Although these data are not sufficient to determine the capacity of the antibodies to neutralize the virus, they support growing evidence of cross-reactivity against H5 HA following influenza vaccinations or infection by other subtypes, indicating that the general population may not be entirely naïve to H5N1 influenza in the case of zoonotic events^31–33^. Similarly, we showed that pre-existing antibodies cross-react with HA from a non-circulating H2N2 strain as well (Fig. 6c).

Serum samples from the BC24 patient demonstrated potent antibody reactivity against the BC24 and MI24 HA proteins (Fig. 6a) and strong cross-reactivity to H1, H2, and H3 HA proteins (Fig. 6b-d). The heightened response to HA is consistent with the timing of the serum collection being at the peak of a typical antibody response to infection. This result suggests that the patient was able to mount a strong humoral response against this BC24 influenza, at least to the HA protein, yet this antibody response was insufficient to mitigate severe disease. In conjunction with evidence of hypercytokinemia^3^, it is also possible that antibody-dependent enhancement contributed to disease severity, but this needs to be investigated further. Nevertheless, our findings show that BC24 HA does not escape targeting by the humoral immune response.

## Conclusion

The cryo-EM structural analyses demonstrate that the E190D and Q226H mutations sequenced from BC24 HA alter the glycan-binding behavior of this HA. Subsequent experiments through *N*-linked glycan microarray, ELISA, and fusion assays in A549 cells support the hypothesis that the BC24 HA has reduced affinity for *N*-linked glycans including both α2,3-linked and α2,6-linked sialosides. We show that the virus likely retains the ability to drive membrane fusion, adding complexity to our understanding of avian influenza’s paths to generating severe disease in humans. Further study will permit a better understanding of how receptor-binding site differences influenced the infectivity of this influenza strain.

## METHODS

### Construct Design and Mutagenesis

VN04 H5 HA: GenBank AAW80717.1; MI24 H5 HA: GenBank XBE32674.1; BC24 H5 HA: GenBank XTJ61994.1; H1 HA: ACP41105.1; H2 HA: AAA43678.1; H3 HA: ACS71642.1 Given the mixed population of HA sequences identified from the patient^3^, the BC24 HA studied here carries both the E190D and Q226H mutations that were most frequently sequenced from patient samples. For structural analysis, we designed soluble HA ectodomain constructs (residues 1-521, H5 sequential numbering) with M36C and G392C (H5 sequential numbering) disulfide-stabilisation mutations^34^ and C-terminal AviTag, thrombin cleavage site, foldon trimerization domain, and 8xHis tag. Since the stem region epitope is distal to the receptor-binding site, the BLI experiments with MEDI8852 were performed using HA carrying an additional Y107F^35^ (H5 sequential numbering) mutation to reduce receptor binding to improve yields and to reduce heterogeneous signal. We expressed these HA with their natural signal peptide sequences from a pcDNA3.1(+) backbone. Plasmids were codon optimized and synthesized at GenScript (Piscataway, NJ, USA). We performed site-directed mutagenesis for the receptor-binding site mutations using the Q5 Site-Directed Mutagenesis kit (New England Biolabs) with validation by whole-plasmid sequencing at Plasmidsaurus (South San Francisco, CA, USA). For fusion assays, we expressed full-length wild-type untagged HA from pcDNA3.1(+) using the natural signal peptide sequences. Plasmids were obtained by custom synthesis at GenScript (Piscataway, NJ, USA).

### HA Soluble Ectodomain Expression and Purification

We expressed HA ectodomains in Expi293F cells (Gibco) for cryo-EM, BLI, and serum ELISA experiments and in GnTI- Expi293F cells (Gibco) for microarray analysis, glycan-binding ELISA experiments, and DLS. We transfected cells at 3×10^6^ cells/mL by complexing HA-encoding plasmids (1.5 μg/mL of culture) with threefold mass excess of linear polyethyleneimine diluted in Expi293 Expression Medium (Gibco). Cells were subsequently incubated at 37°C in 8% CO_2_ at 130 rpm (25 mm orbital radius) with addition of 2.2 mM valproic acid (100X in phosphate buffered saline (PBS) pH 7.4) 1 day post-transfection. 5 days post-transfection, we harvested media supernatants containing HA ectodomain by centrifugation at 500×g then clarified them by centrifugation at 15 000×g and filtration through a 0.45 μm pore size PES membrane.

We purified HA from the clarified supernatants on an ÄKTA pure system (GE Healthcare) at 4°C by nickel affinity chromatography (loading onto a 5 mL HisTrap HP (Cytiva); washing with 20 mM Tris pH 8, 500 mM NaCl, 20 mM imidazole; and elution in 20 mM Tris pH 8, 500 mM NaCl, 500 mM imidazole). For microarray, glycan-binding ELISA, and DLS experiments, we buffer exchanged affinity-purified product with Amicon Ultra centrifugal filters (Millipore), 100 kDa molecular-weight cutoff (4 mL) to TBS (20 mM Tris pH 8, 150 mM NaCl). For cryo-EM, BLI, DLS (where specified), and serum ELISA experiments, we further purified samples by size-exclusion chromatography (SEC; Superdex 200 Increase 30/100 GL (Cytiva), eluted with TBS) before concentration by centrifugal filtration to 1.7-2.0 mg/mL.

### Cryo-Electron Microscopy

We applied 1.8 μL of purified HA ectodomains at 10°C, >98% humidity onto Quantifoil R1.2/1.3, 200 mesh, Cu holey carbon grids glow discharged for 20 s using a Pelco easiGlow. Using a Vitrobot Mark IV (Thermo Fisher Scientific), we then blotted grids for 12 s at-10 relative blot force and plunge-froze samples in liquid ethane. We collected micrographs using a Titan Krios G4 transmission electron microscope (Thermo Fisher Scientific) at 300 kV with a Selectris X imaging filter set to 10 eV slit width and Falcon 4 direct electron detector in electron event registration mode. We collected movies at 165 000x magnification over a defocus range of-1.0 to-2.0 μm with a total dose of 40.0 e^-^/Å^2^.

We performed data processing using CryoSPARC v4.6, with the workflow provided in Supplementary Fig. 1-3. In brief, following patch motion correction and CTF estimation, we performed blob particle picking. After iterative rounds of heterogeneous refinement, we generated the final reconstruction with non-uniform refinement with C3 symmetry imposed. We fit an initial model of a F_ab_-complexed VN04 HA ectodomain crystal structure into the maps using UCSF ChimeraX^36^. We then subjected the model to several cycles of real-space refinement in Phenix v1.21.1 and manual adjustment on WinCoot v0.9.8^37,38^. Model statistics are available in Supplementary Data 1. We introduced mutations as needed using WinCoot. Figures were prepared using UCSF ChimeraX.

### MEDI8852 F_ab_ Preparation

We expressed MEDI8852 F_ab_ (UniRef100_UPI0008141F01, UniRef100_UPI0008141F29) from pcDNA3.1(+) backbones, with the heavy chain carrying a C-terminal 6xHis tag. Plasmids were synthesised by GenScript. We expressed MEDI8852 F_ab_ in Expi293F cells in the same manner as the soluble HA ectodomains, transfecting with 1 μg each of heavy-and light-chain-encoding plasmid per mL of culture. We purified F_ab_s by nickel affinity chromatography and size-exclusion chromatography in the same manner as the soluble HA ectodomains.

### *N*-Glycan Microarray

Microarray experiments using the Z Biotech N-glycan array kit (10602-16K) and signal processing were performed by Creative Proteomics (Shirley, NY, USA). His-tagged HA ectodomains (20 μg/mL) were precomplexed in glycan array assay buffer (GAAB; Z Biotech) at a 10:1:1 mass ratio with rabbit anti-6xHis antibody (Invitrogen PA1-983B) and Cy3-conjugated anti-rabbit IgG (Life Technologies A10520) for 1 h on ice. The microarray was blocked for 30 min at room temperature in glycan array blocking buffer (Z Biotech). 100 μL of precomplex solution was added to each subarray and incubated for 1 h at room temperature prior to washing in GAAB then water. The microarray slide was then scanned at 532 nm. Analysis was performed on Mapix (Innopsys) to subtract background and signal from dedicated negative control spots without printed glycan. Fig. 3 presents the data reordered to categorise glycans by terminal linkage type, from left to right N1, N7, N13, N19, N25, N31, N37, N41, N46, N53, N57, N62, N82, N87, N91, N96, N102, N106, N111, N4, N10, N16, N22, N28, N34, N40, N44, N47, N50, N56, N60, N63, N85, N90, N94, N97, N100, N105, N109, N112, N45, N49, N61, N65, N95, N99, N110, N114, N2, N8, N14, N20, N26, N32, N38, N42, N48, N54, N58, N64, N83, N88, N92, N98, N103, N107, N113, N3, N5, N6, N9, N11, N12, N15, N17, N18, N21, N23, N24, N27, N29, N30, N33, N35, N36, N39, N43, N51, N52, N55, N59, N66, N67, N68, N69, N70, N71, N72, N73, N74, N75, N76, N77, N78, N79, N80, N81, N84, N86, N89, N93, N101, N101, N108, according to the Z Biotech labels (structures and labels provided Supplementary Data 2).

### Dynamic Light Scattering

We analysed using a Prometheus Panta (NanoTemper) the protein preparations used for *N*-glycan microarray analysis as well positive and negative comparisons. For a more purified preparation, BC24 HA ectodomain from GnTI- Expi293F cells further purified by SEC was included. For an aggregated comparison, we subjected a portion of the BC24 HA sample (not purified by SEC) to four rounds of freezing and thawing. 0.2 mg/mL HA ectodomains were assayed on a Prometheus Panta in technical triplicate. The “Size Analysis” dynamic light scattering (DLS) procedure was run in high sensitivity mode at 25°C. Data were processed automatically by Panta software, and figures were generated with the same software.

### Glycan-Binding ELISA Experiments

We immobilised biotin-conjugated biantennary glycans (Sussex Research BT000040, BT000055, BT000050, and BT000060) on Reacti-Bind streptavidin-coated 384-well plates (Pierce) at 4°C for 16 h in PBS pH 7.4. Wells were then blocked in 6 mM d-desthiobiotin in PBS pH 7.4 for 1.5 h at room temperature. We concentrated preparations of soluble HA ectodomains to 1.5-3.5 mg/mL prior to precomplexing for 1 h at room temperature with peroxidase-conjugated rabbit anti-His tag IgG (Jackson ImmunoResearch 300-035-240) at a 2:1 molar ratio of HA trimer to antibody. We prepared this precomplex solution in TBS supplemented with 1% (w/v) bovine serum albumin (BSA). We then added HA-antibody complexes at the desired dilutions in BSA-supplemented TBS to the plate for incubation at room temperature for 2 h. Following washing five times in PBS pH 7.4 with 0.05% Tween-20 (PBS-T), we developed signal by addition of Pierce 1-Step Turbo TMB-ELISA substrate for 13 min at room temperature before quenching by addition of an equal volume of 2 M sulfuric acid. We then immediately measured absorbance at 450 nm using a Varioskan LUX (Thermo Scientific). We used GraphPad Prism 10 for all data analysis and binding curve generation. Using the “Sigmoidal dose-response (variable slope)” method, we fit curves for data with log_10_-transformed concentration values then averaged logEC_50_ values across biological replicates. Error bars are standard deviation. We produced glycan schematics using GlycoGlyph then adjusted them on Inkscape^39^.

### Cell-Based HA Fusion Assay

We seeded 2×10^4^ A549 cells (ATCC) onto transparent-bottomed, black-walled 96-well plates (Greiner Bio-One) in Dulbecco’s modified Eagle’s medium (DMEM) supplemented with 10% heat-inactivated fetal bovine serum and penicillin-streptomycin. We incubated cultures at 37°C in 5% CO_2_. 16 h post-seeding, we transfected cells following the Lipofectamine 3000 protocol (Invitrogen). For each technical replicate, we transfected one group of cells to express GFP-HiBiT (Addgene 162589), and the other to express both the full-length HA of interest and mCherry-lgBiT (Addgene 199715). After an additional 24 h, we detached HA-transfected cells by 8 min incubation in 0.025% trypsin-EDTA at 37°C and transferred them onto the GFP-HiBiT-transfected cells to incubate for 24 h. We activated HA by replacing the media with 90 mM sodium citrate pH 4.5, 150 mM NaCl, prewarmed to 37°C, for 3 min, then immediately washed the cells in supplemented DMEM and let them recover in a second change of media for 6 h. We measured reconstituted NanoBiT activity as a proxy of fusion using the Nano-Glo Live Cell Assay System (Promega), recording luminescence with 1 s integration time 5 min after addition of substrate using a Varioskan LUX at 37°C. We determined significance values on GraphPad Prism 10 by two-tailed t-test with Welch’s correction, with parameters provided in Supplementary Table 1.

We immediately fixed cells by incubation for 20 min in 4% paraformaldehyde (Electron Microscopy Sciences) diluted in PBS pH 7.4, with a wash in PBS both before and after fixation. Following incubation in 1 μg/mL 4′,6-diamidino-2-phenylindole (DAPI) for 10 min, we washed wells five times in PBS then stored them in ProLong Glass Antifade (Invitrogen) before imaging on a Zeiss LSM 880 confocal microscope. We acquired a z-stack of five 2 μm-thick optical sections, and 2D projections are presented.

We performed inhibition assays in the same manner except supplementing media with the indicated concentration of MEDI8852 F_ab_ 1 h prior to acidification, during which time cells remained at 37°C with 5% CO_2_. We performed data analysis on GraphPad Prism 10 in the same manner as for glycan-binding ELISA experiments except that data were first normalised within the range of signal for each technical replicate before merging and calculating IC_50_ values.

### Western Blotting to Detect HA Expression

We harvested fusion assay A549 cells co-transfected with mCherry-lgBiT and the indicated H5 HA (Supplementary Fig. 7) by addition of 0.2 M Tris pH 6.5, 8% (w/v) sodium dodecyl sulfate (SDS), 4.3 M glycerol, 6 mM bromophenol blue, 0.4 M dithiothreitol. We then heated harvested samples at 95°C for 10 min and loaded one third of the sample onto a 4-20% gradient Mini-PROTEAN TGX gel (Bio-Rad) to be run at 140 V submerged in Tris/glycine/SDS running buffer (Bio-Rad). Following blotting onto a nitrocellulose membrane at 100 V for 75 min in 4°C 25 mM Tris, 192 mM glycine, 10% (v/v) methanol, we validated transfer using Ponceau S. We washed the membrane several times in water then blocked it for 10 min at room temperature in EveryBlot Blocking Buffer (Bio-Rad). We incubated the blot in 1:2000 rabbit anti-HA mAb (Sino Biological 86001-RM01) and 1:1500 BA3R mouse anti-β-actin loading control mAb (Invitrogen) diluted in EveryBlot Blocking Buffer for 18 h at 4°C. Following washing three times for ten minutes in PBS-T, we then incubated the blot in 1:2000 peroxidase-conjugated anti-mouse IgG (Jackson ImmunoResearch 715-035-150) and 1:2000 anti-rabbit IgG antibodies (Promega W401B). After washing five times for ten minutes each in PBS-T, we developed signal using Clarity Western ECL Substrate (Bio-Rad) and imaged the blot on a ChemiDoc MP imaging system (Bio-Rad) with both colorimetric (to validate band sizes against the stained protein standards) and chemiluminescent channels.

### Biolayer Interferometry to Assay MEDI8852 F_ab_:HA Binding

We performed all experiments on a Gator Prime system (Gator Bio) set to 30°C and in BLI buffer (TBS supplemented with 0.02% Tween-20). We equilibrated anti-human FAB probes (Gator Bio) for 600 s at 400 rpm in BLI buffer before a 120 s baseline step at 1000 rpm. We loaded MEDI8852 F_ab_ diluted to 100 nM onto probes for 40 s with 400 rpm shaking, reaching ∼1 nm loading shift, followed by another 120 s baseline at 1000 rpm. We carried out both the HA ectodomain association and dissociation steps for 1200 s at 1000 rpm. HA:HA interactions may result in inflated K_D_ especially for the VN04 and MI24 HA proteins which have higher glycan affinities, so we sought to limit this effect by using Y98F HA mutants with reduced sialoside affinities^35^.

We used GatorOne software (Gator Bio) for data processing and analysis. Processing included y-axis alignment of the association phase, inter-step correction around the dissociation phase, and double reference subtraction. Processed sensograms are presented with Savitzky-Golay filtering. We determined kinetics and affinity parameters presented in Supplementary Table 2 by fitting curves using a 2:1 global binding model with R_max_ considered unlinked. We recreated sensograms using the seaborn version 0.13.0 library for Python.

### Patient Sample Collection

Sera from BC24 patient and healthy age-matched controls were obtained at the British Columbia Children’s Hospital and stored at their BioBank according to ethics approval from the UBC C&W Research Ethics Board (protocol H23-02299). Use of the collected samples for research was approved by the UBC Clinical Research Ethics Board (H24-03937).

### ELISA to Assay Serum Reactivity to Clade 2.3.4.4b HA

BC24 and MI24 HA ectodomains were coated on a 96-well Nunc MaxiSorp plate (Thermo Scientific) by incubation of 2 μg/mL HA overnight at 4°C. Wells were subsequently washed three times in PBS-T before blocking for 2 h at room temperature in PBS supplemented with 1% bovine serum albumin (blocking buffer). Wells were then incubated overnight at 4°C in serum samples serially diluted in blocking buffer, washed, and subsequently incubated in 1:5000 peroxidase-conjugated goat anti-human IgG (Jackson ImmunoResearch) for 1 h at room temperature before washing three times in PBS-T. We developed the plates with TMB substrate and measured signal at OD450nm.

## ACKNOWLEDGEMENTS

This work was supported by an award from the Canada Biomedical Research Fund (CBRF) for the PROGENITER pandemic preparedness project, and by awards to S.S. from a Canada Excellence Research Chair Award and the VGH Foundation. J.H.N. was supported by a University of British Columbia (UBC) Four Year Doctoral Fellowship and UBC President’s Academic Excellence Initiative PhD Award. We would like to thank Dima Lyubashenko and Dhiraj Mannar for helpful contributions, and Stefan Wendt, Ada Lin, and the Nygaard Lab for use and guidance of their confocal microscope. We would like to thank Tatiana Lau in the Tokuyama lab for technical assistance. We gratefully acknowledge the work and staff of the BC Children’s Hospital BioBank for their assistance with sample acquisition. We also wish to thank all members of the Subramaniam laboratory and the PROGENITER team for helpful suggestions and comments throughout the course of this work.

## SUPPLEMENTARY INFORMATION

**Supplementary Figure 1.**
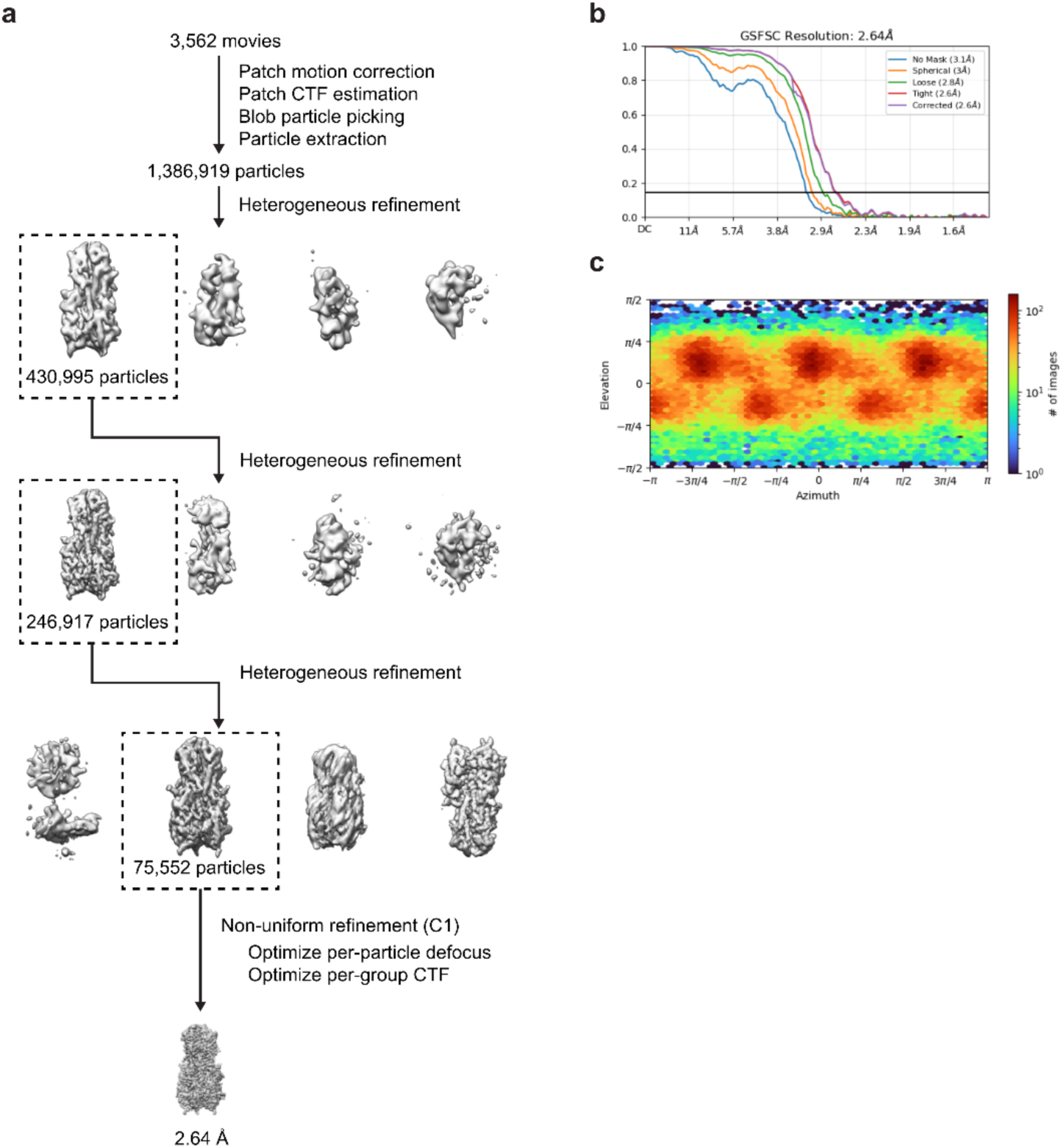
Cryo-EM data processing for VN04 HA ectodomain. **a.** Steps in cryo-EM processing. **b.** Fourier shell correlation curves. **c.** Viewing direction distribution plot.

**Supplementary Figure 2.**
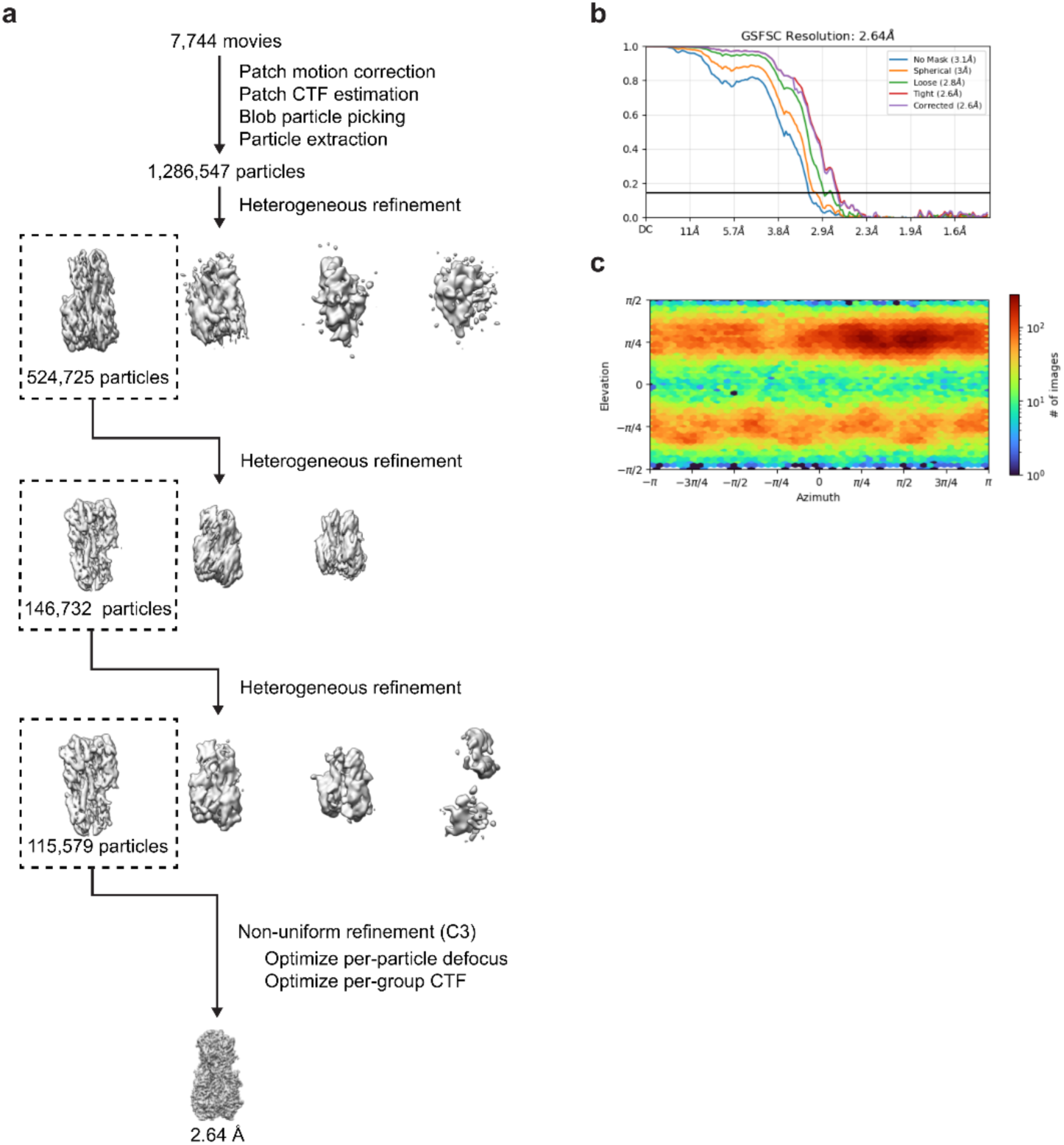
Cryo-EM data processing for MI24 HA ectodomain. **a.** Steps in cryo-EM processing. **b.** Fourier shell correlation curves. **c.** Viewing direction distribution plot.

**Supplementary Figure 3.**
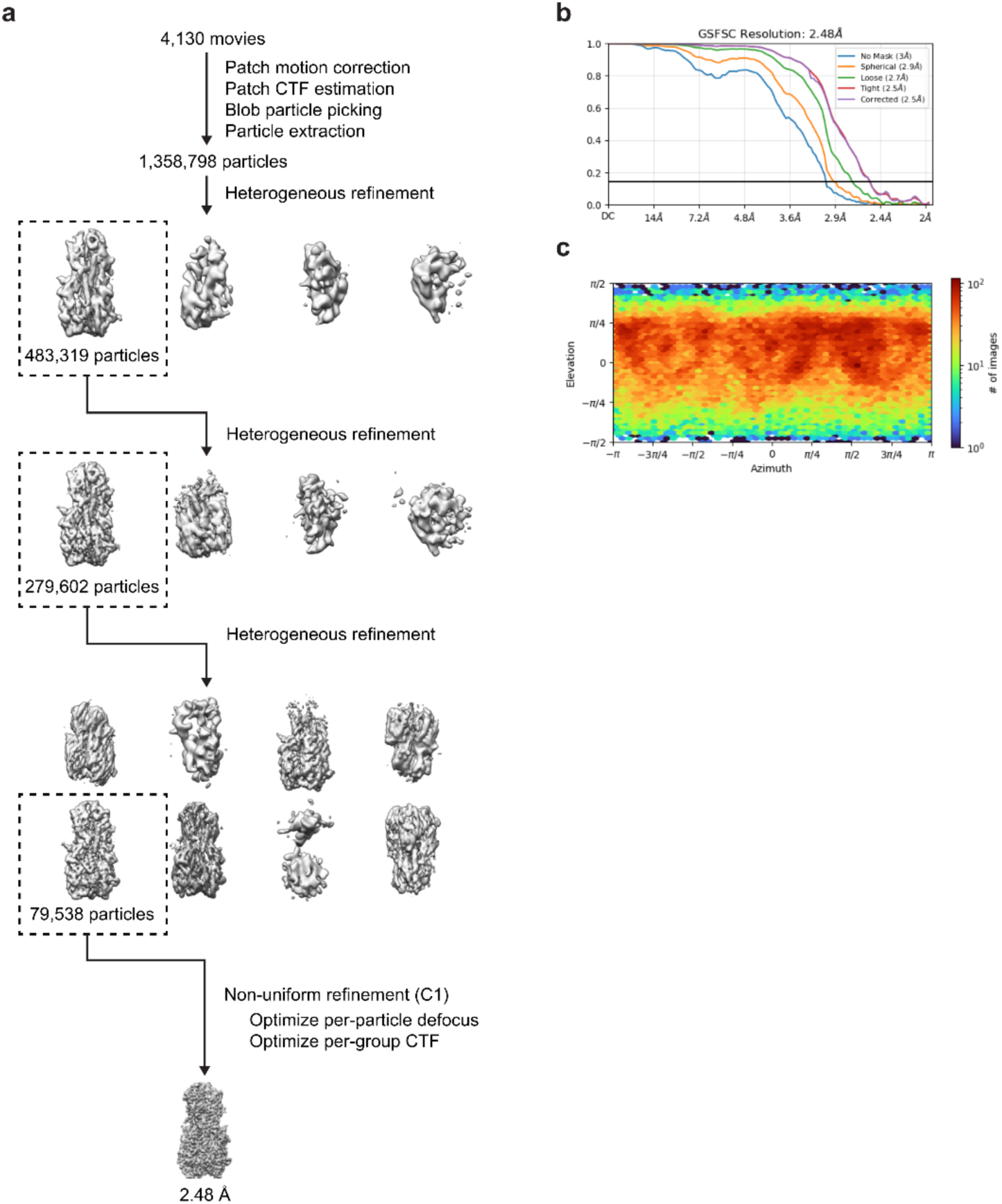
Cryo-EM data processing for BC24 HA ectodomain. **a.** Steps in cryo-EM processing. **b.** Fourier shell correlation curves. **c.** Viewing direction distribution plot.

**Supplementary Figure 4.**
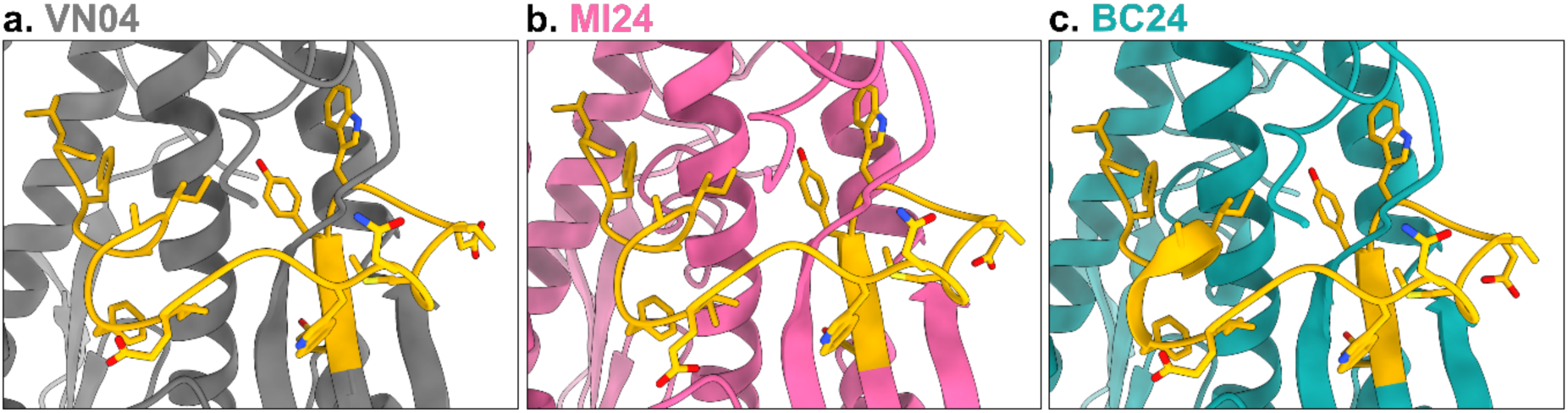
Fusion peptide in pre-fusion state. From HA ectodomains expressed in regular Expi293F cells, zoomed-in views of cryo-EM-derived models are presented for the fusion peptide (yellow) of **a.** VN04, **b.** MI24, and **c.** BC24 HA proteins.

**Supplementary Figure 5.**
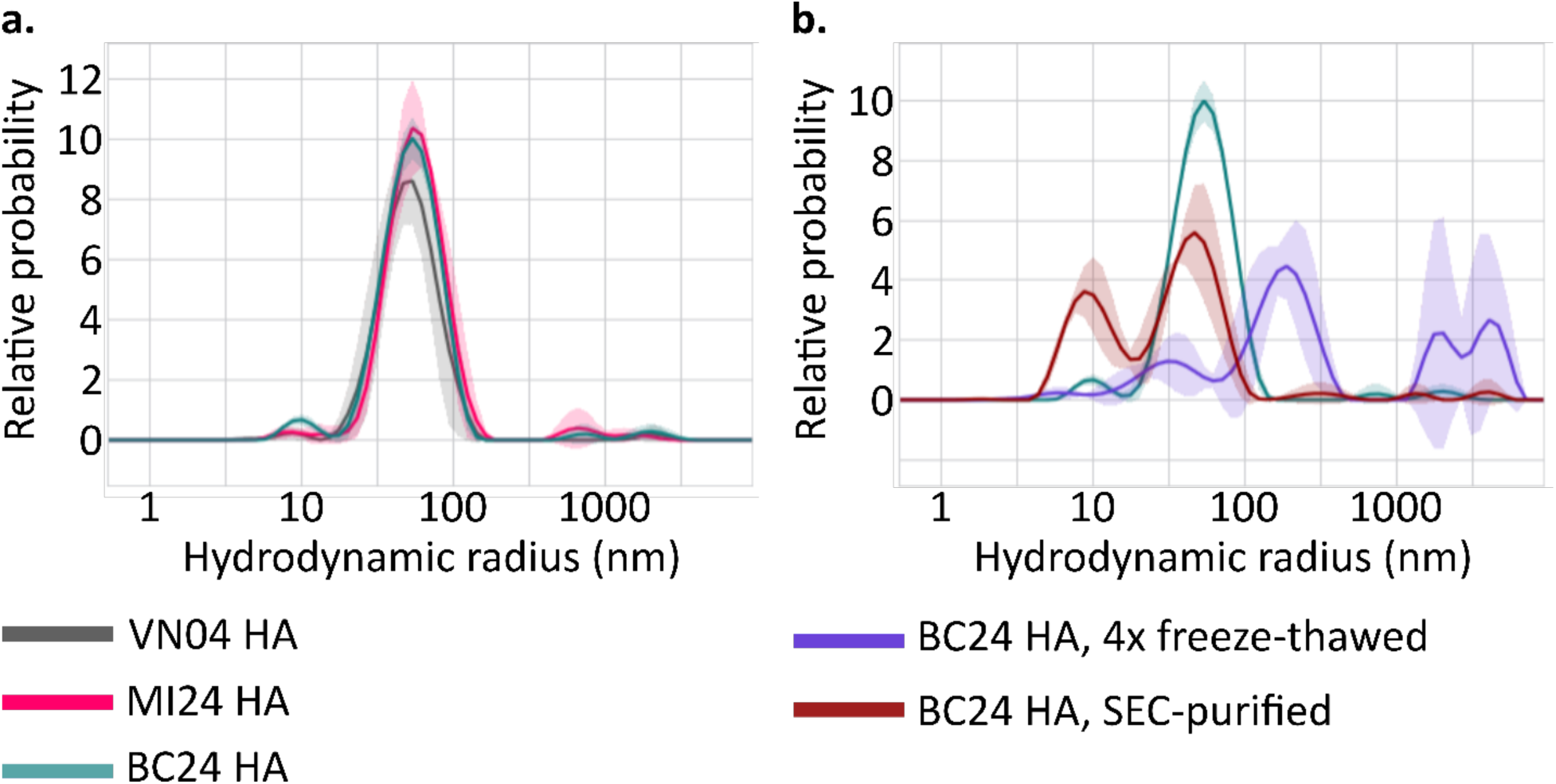
Dynamic light scattering demonstrates comparable protein quality in microarray analysis. HA ectodomain preparations used in microarray experiments were analysed by DLS at 25°C. **a.** Size distribution of the VN04 (grey), MI24 (pink), and BC24 (teal) HA ectodomain preparations. Major peaks at hydrodynamic radius ≈ 50 nm demonstrate HA:HA association of intact trimers, whereas the peaks at ∼9 nm are assigned as individual HA trimers. **b.** Comparison of the BC24 HA preparations used for microarray analysis (teal), a preparation additionally purified by SEC (rust), and one subjected to four freeze-thaw cycles to promote aggregation (purple). Shaded regions are standard deviation.

**Supplementary Figure 6.**
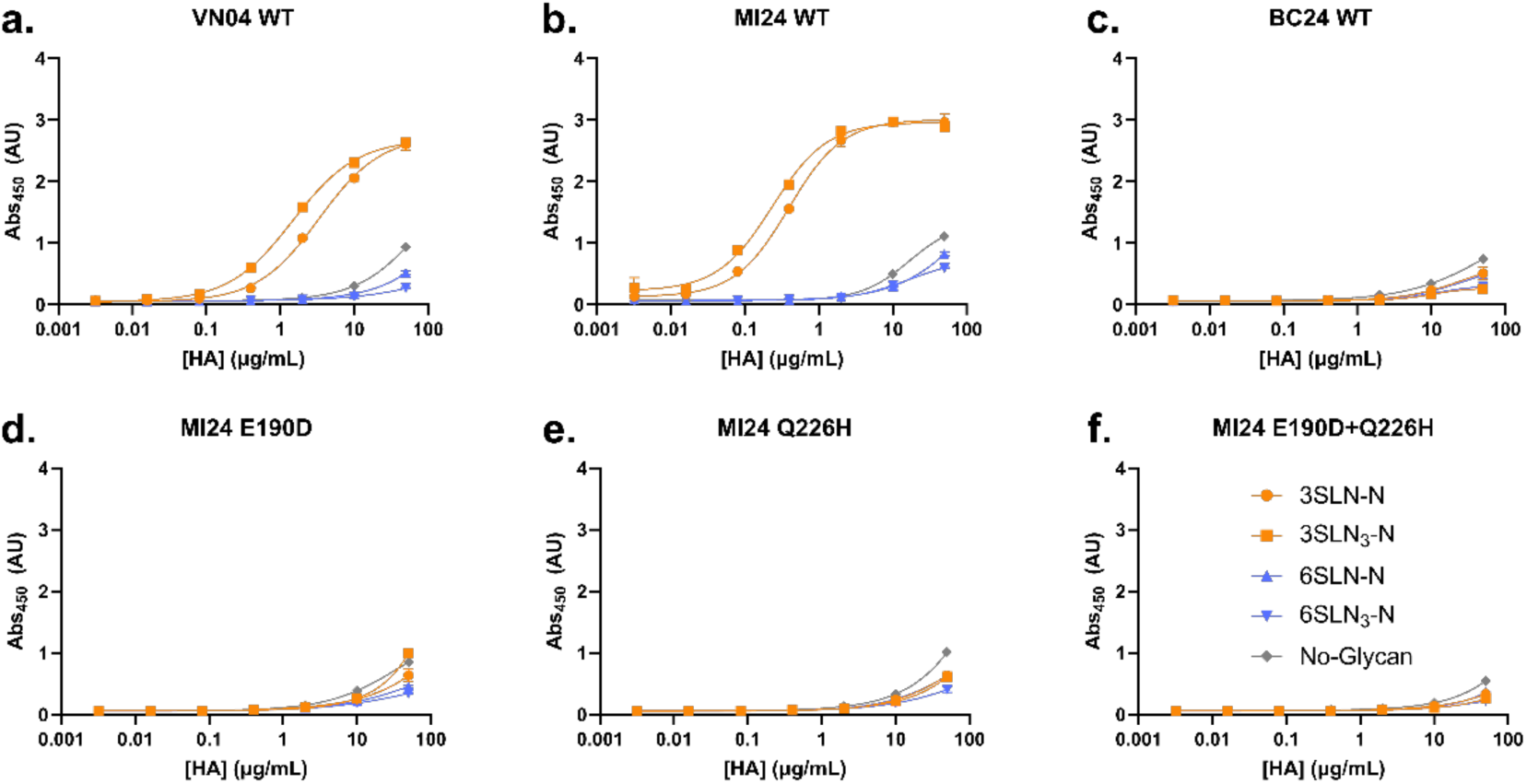
Biological replicate of glycan-binding ELISA with H5 HA proteins. Using separately prepared protein, ELISA curves were generated probing binding to biantennary α2,3- and α2,6-linked sialosides by HA variants **a.** VN04 wild-type, **b.** MI24 wild-type, **c.** BC24 wild-type, **d.** MI24 E190D mutant, **e.** MI24 Q226H mutant, and **f.** MI24 E190D+Q226H double mutant. Error bars are standard deviation (n=3).

**Supplementary Figure 7.**
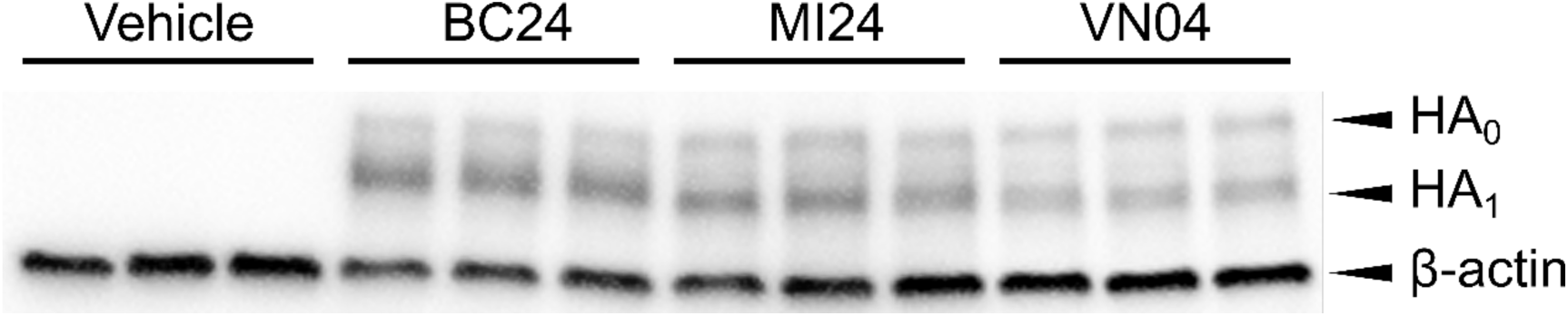
Differences in fusion efficiency are not due to expression differences. A549 cells co-transfected with mCherry-lgBiT and the indicated HA alongside fusion assay samples were analysed in triplicate by western blotting, probing for HA and β-actin, with corresponding bands labelled to the right of the blot. Secondary antibodies are peroxidase-conjugated, and the chemiluminescent scan is presented.

**Supplementary Figure 8.**
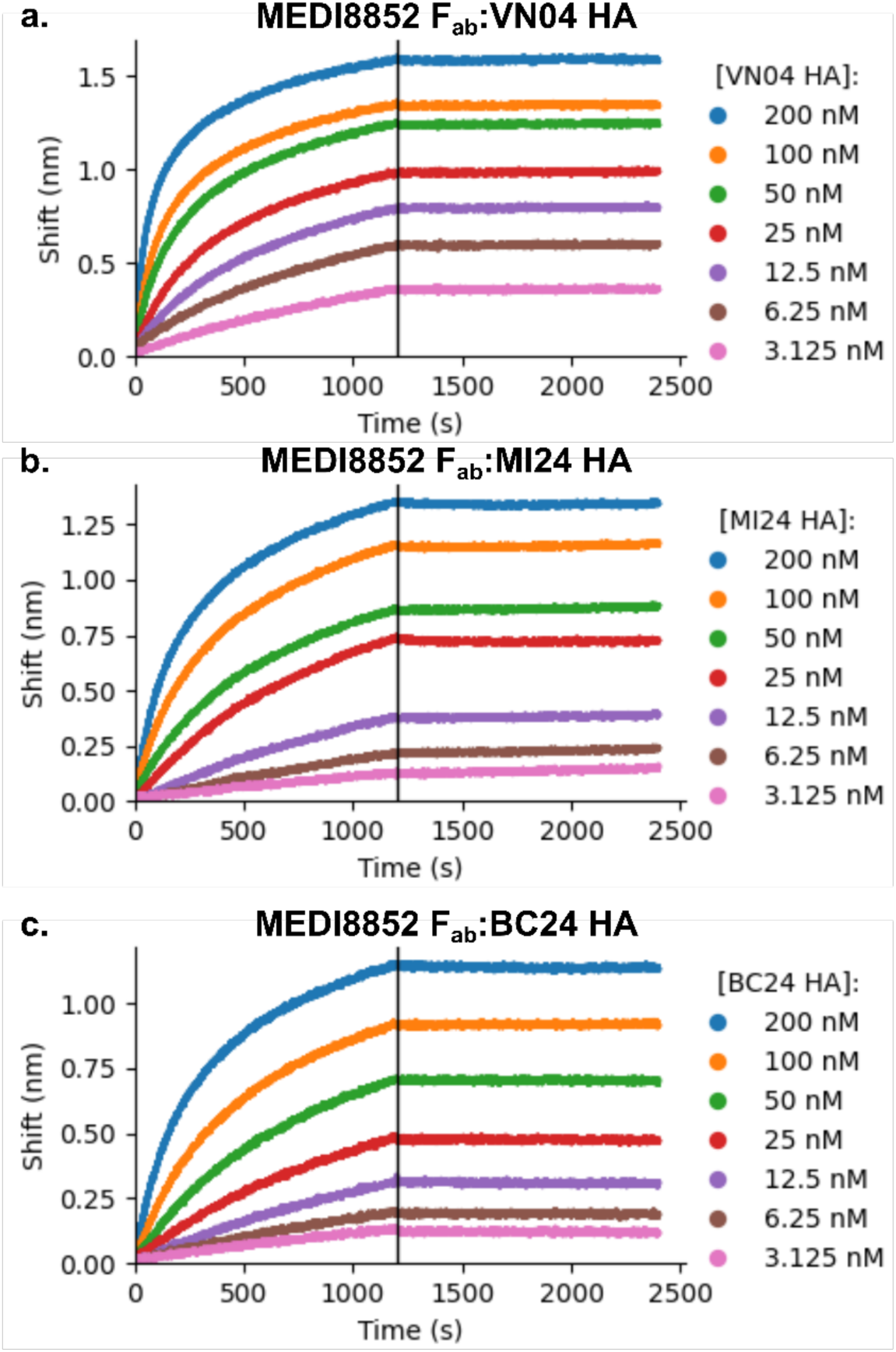
BLI sensograms for MEDI8852:HA-binding. Sensograms after double reference subtraction for MEDI8852 F_ab_ binding to **a.** VN04, **b.** MI24, and **c.** BC24 H5 HA proteins. Vertical marker indicates switch from association to dissociation phase.

---

**Supplementary Table 1.** Statistics parameters for fusion assay data, HA-transfected compared to mock-transfected control.

| HA | t-Value | Degrees of Freedom | p-Value |
| --- | --- | --- | --- |
| VN04 | 13.07 | 7.290 | <0.0001 |
| MI24 | 6.346 | 5.289 | 0.0012 |
| BC24 | 5.174 | 8.712 | 0.0006 |

**Supplementary Table 2.** Summary of BLI data demonstrating MEDI8852 binding to all H5 HA studied.

| HA | $k_{\text{off}}$ (s<sup>-1</sup>) | $k_{\text{on}}$ (M<sup>-1</sup>s<sup>-1</sup>) | K<sub>D</sub> (nM) |
| --- | --- | --- | --- |
| VN04 | 5.69×10<sup>-6</sup> ±<br>4.48×10<sup>-7</sup> | 1.49×10<sup>4</sup> ±<br>3.73×10<sup>1</sup> | 0.383 |
| MI24 | 8.18×10<sup>-6</sup> ±<br>1.54×10<sup>-7</sup> | 9.65×10<sup>3</sup> ±<br>1.61×10<sup>1</sup> | 0.848 |
| BC24 | 1.43×10<sup>-6</sup> ±<br>1.17×10<sup>-7</sup> | 8.59×10<sup>3</sup> ±<br>1.92×10<sup>1</sup> | 0.167 |

**Supplementary Table 3.** Patient* data for serum reactivity ELISA.

| Sample | Gender | Collection Timepoint** |
| --- | --- | --- |
| BC24-1 | Female | Timepoint 1 |
| BC24-2 | Female | Timepoint 2 |
| BC24-3 | Female | Timepoint 3 |
| BC24-4 | Female | Timepoint 4 |
| Healthy-1 | Male | NA |
| Healthy-2 | Male | NA |
| Healthy-3 | Female | NA |
| Healthy-4 | Female | NA |

\*The ages of the healthy donors and the patient were all within the range of 13-15 years.
\*\*The samples were collected 1-4 days after a negative PCR test for the virus in the serum and estimated to be 2-3 weeks after the initial infection.

**Supplementary Data 1.**
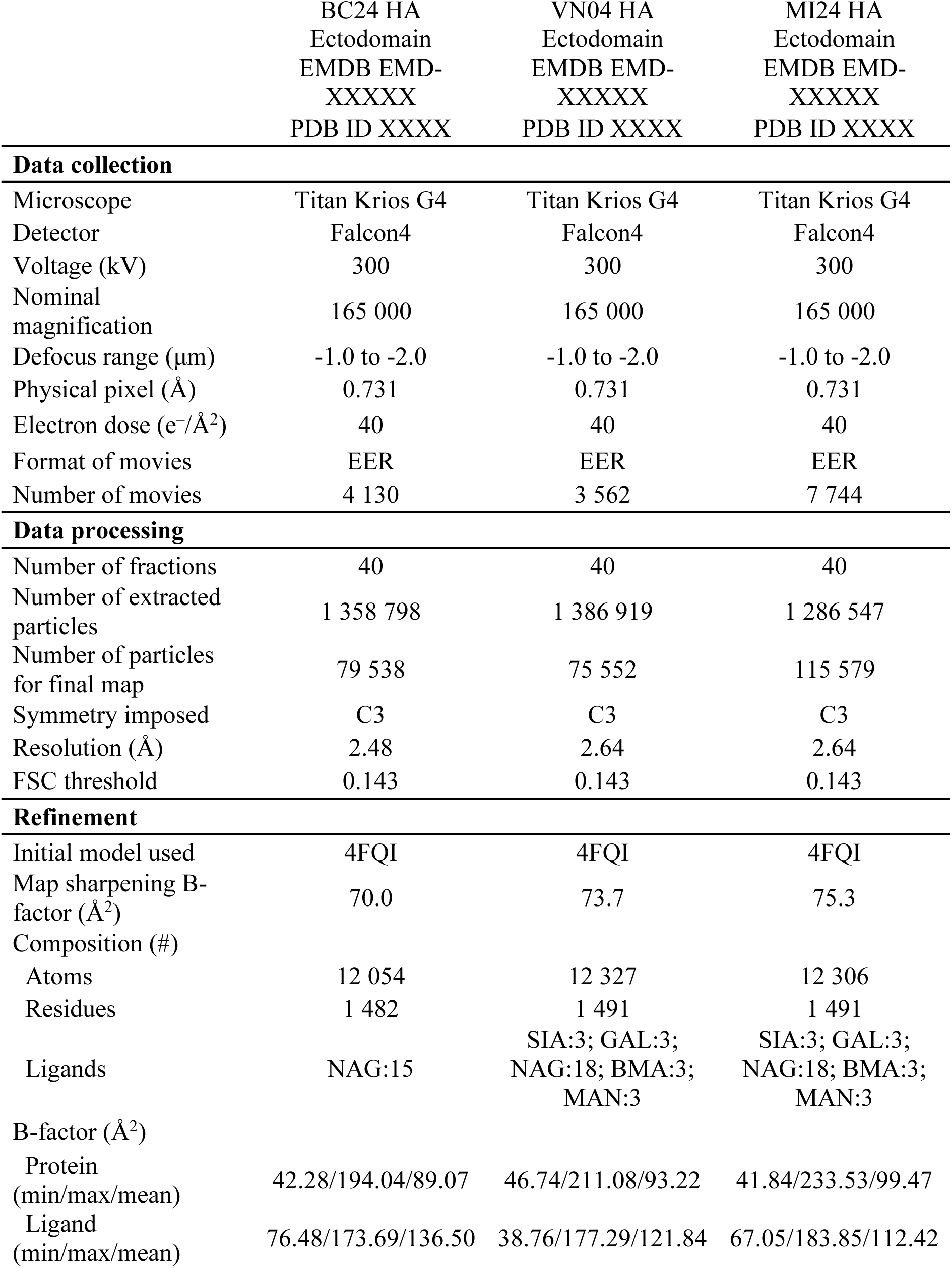
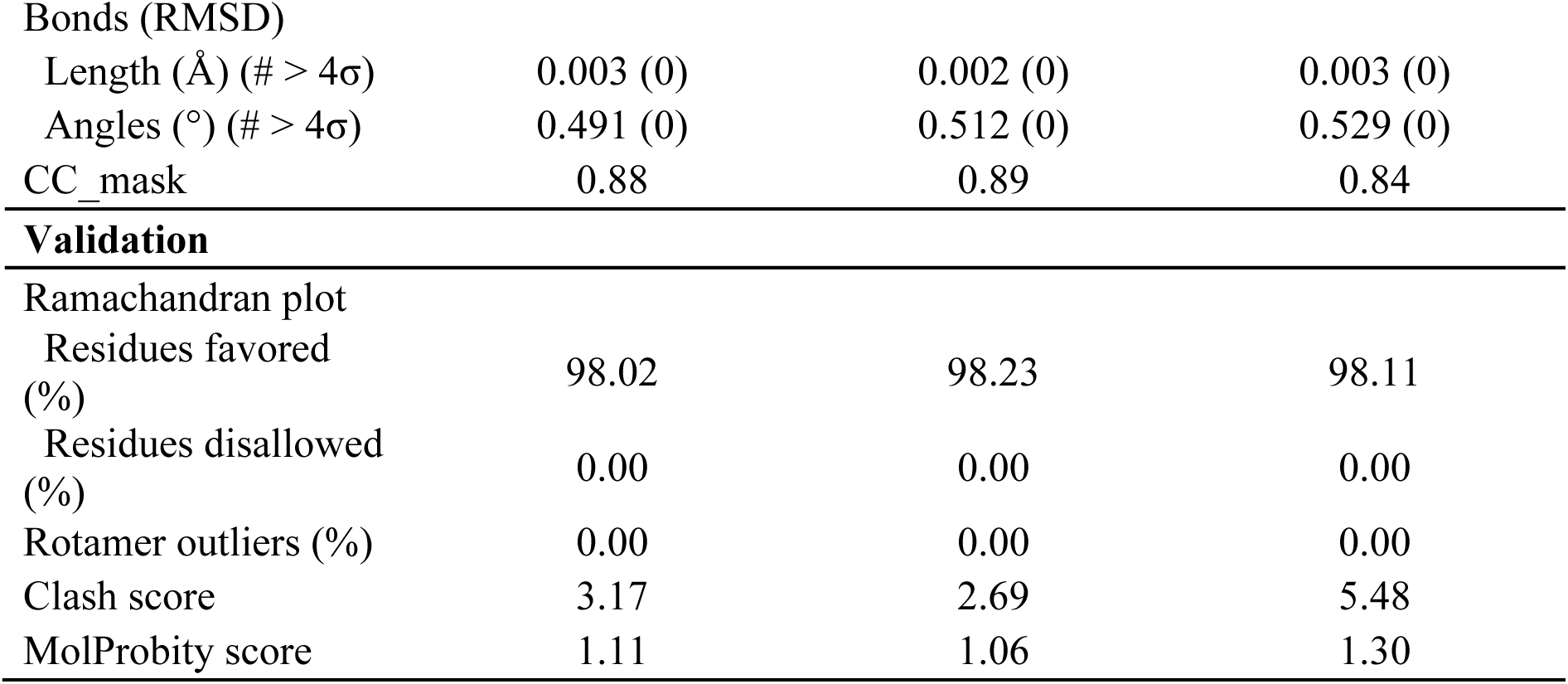
Data collection, data processing, and model validation statistics for reported structures.

|  | BC24 HA<br>Ectodomain<br>EMDB EMD-XXXXX<br>PDB ID XXXX | VN04 HA<br>Ectodomain<br>EMDB EMD-XXXXX<br>PDB ID XXXX | MI24 HA<br>Ectodomain<br>EMDB EMD-XXXXX<br>PDB ID XXXX |
| --- | --- | --- | --- |
| **Data collection** |  |  |  |
| Microscope | Titan Krios G4 | Titan Krios G4 | Titan Krios G4 |
| Detector | Falcon4 | Falcon4 | Falcon4 |
| Voltage (kV) | 300 | 300 | 300 |
| Nominal magnification | 165 000 | 165 000 | 165 000 |
| Defocus range ($\mu\text{m}$) | -1.0 to -2.0 | -1.0 to -2.0 | -1.0 to -2.0 |
| Physical pixel ($\text{\AA}$) | 0.731 | 0.731 | 0.731 |
| Electron dose ($\text{e}^-/\text{\AA}^2$) | 40 | 40 | 40 |
| Format of movies | EER | EER | EER |
| Number of movies | 4 130 | 3 562 | 7 744 |
| **Data processing** |  |  |  |
| Number of fractions | 40 | 40 | 40 |
| Number of extracted particles | 1 358 798 | 1 386 919 | 1 286 547 |
| Number of particles for final map | 79 538 | 75 552 | 115 579 |
| Symmetry imposed | C3 | C3 | C3 |
| Resolution ($\text{\AA}$) | 2.48 | 2.64 | 2.64 |
| FSC threshold | 0.143 | 0.143 | 0.143 |
| **Refinement** |  |  |  |
| Initial model used | 4FQI | 4FQI | 4FQI |
| Map sharpening B-factor ($\text{\AA}^2$) | 70.0 | 73.7 | 75.3 |
| Composition (#) |  |  |  |
| Atoms | 12 054 | 12 327 | 12 306 |
| Residues | 1 482 | 1 491 | 1 491 |
| Ligands | NAG:15 | SIA:3; GAL:3;<br>NAG:18; BMA:3;<br>MAN:3 | SIA:3; GAL:3;<br>NAG:18; BMA:3;<br>MAN:3 |
| B-factor ($\text{\AA}^2$) | | | |
| Protein (min/max/mean) | 42.28/194.04/89.07 | 46.74/211.08/93.22 | 41.84/233.53/99.47 |
| Ligand (min/max/mean) | 76.48/173.69/136.50 | 38.76/177.29/121.84 | 67.05/183.85/112.42 |
| Bonds (RMSD) |  |  |  |
| Length (Å) (# > 4σ) | 0.003 (0) | 0.002 (0) | 0.003 (0) |
| Angles (°) (# > 4σ) | 0.491 (0) | 0.512 (0) | 0.529 (0) |
| CC_mask | 0.88 | 0.89 | 0.84 |
| **Validation** |  |  |  |
| Ramachandran plot |  |  |  |
| Residues favored (%) | 98.02 | 98.23 | 98.11 |
| Residues disallowed (%) | 0.00 | 0.00 | 0.00 |
| Rotamer outliers (%) | 0.00 | 0.00 | 0.00 |
| Clash score | 3.17 | 2.69 | 5.48 |
| MolProbity score | 1.11 | 1.06 | 1.30 |

**Supplementary Data 2.**
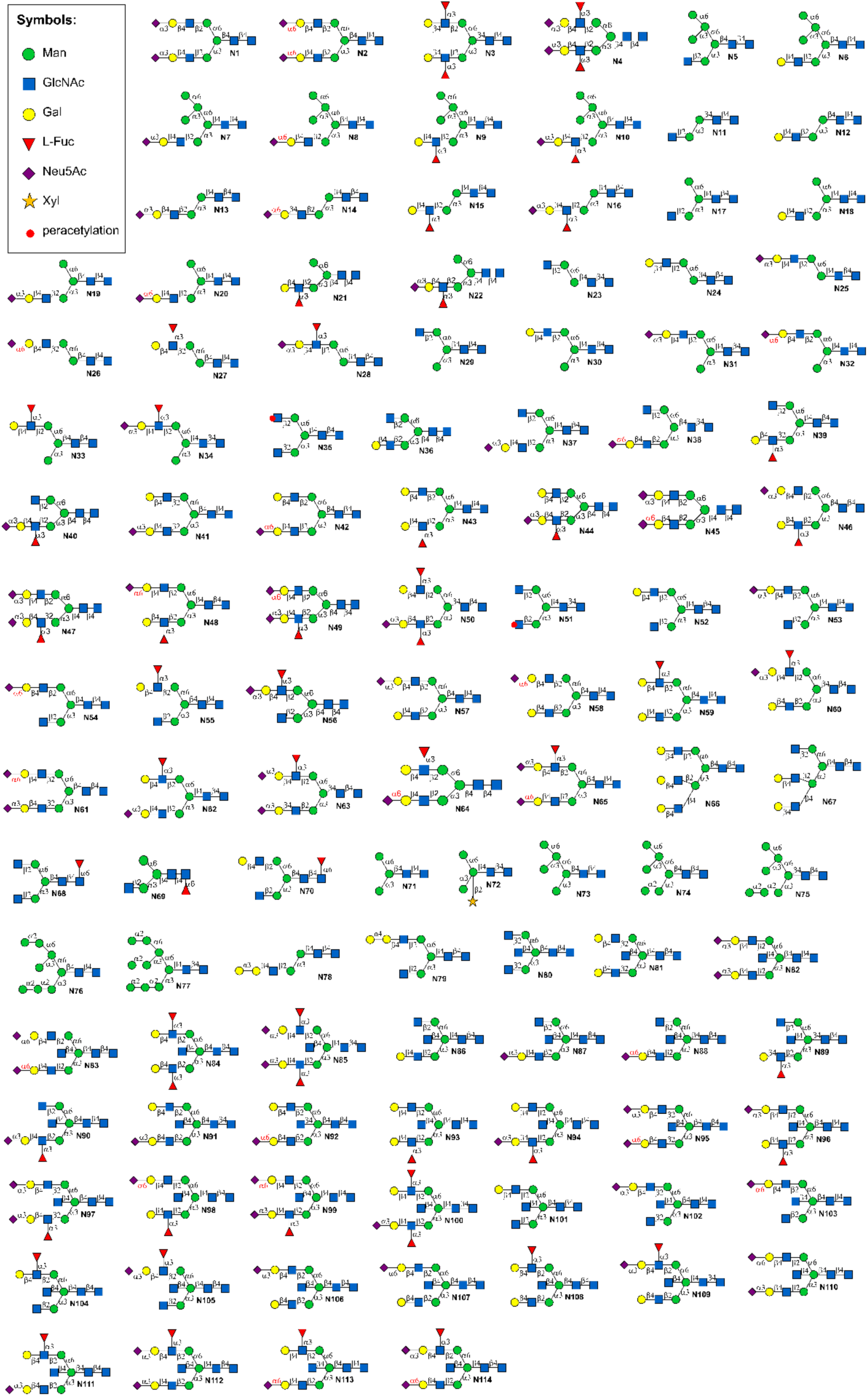
Structures of *N*-glycans assayed by microarray analysis. . Reproduced with permission from Z Biotech.

## REFERENCES

1 Himsworth, C. et al. Highly Pathogenic Avian Influenza A(H5N1) in Wild Birds and a Human, British Columbia, Canada, 2024. Emerg. Infect. Dis. 31, 1216 (2025). 10.3201/eid3106.241862

2 World Health Organization. Cumulative number of confirmed human cases for avian influenza A(H5N1) reported to WHO, 2003-2025, 20 January 2025. Cumulative number of confirmed human cases for avian influenza A(H5N1) reported to WHO, 2003-2025, 20 January 2025, https://cdn.who.int/media/docs/default-source/2021-dha-docs/cumulative-number-of-confirmed-human-cases-for-avian-influenza-a(h2025n2021)-reported-to-who--2003-2025.pdf (2025).

3 Jassem, A. N. et al. Critical Illness in an Adolescent with Influenza A(H5N1) Virus Infection. N. Engl. J. Med. 392, 927–929 (2024). 10.1056/NEJMc2415890

4 Nicholls, J. M., Bourne, A. J., Chen, H., Guan, Y. & Peiris, J. S. Sialic acid receptor detection in the human respiratory tract: evidence for widespread distribution of potential binding sites for human and avian influenza viruses. Respir. Res. 8, 73 (2007). 10.1186/1465-9921-8-73

5 Dadonaite, B. et al. Deep mutational scanning of H5 hemagglutinin to inform influenza virus surveillance. PLoS Biol. 22, e3002916 (2024). 10.1371/journal.pbio.3002916

6 Glaser, L. et al. A single amino acid substitution in 1918 influenza virus hemagglutinin changes receptor binding specificity. J. Virol. 79, 11533–11536 (2005). 10.1128/jvi.79.17.11533-11536.2005

7 Stevens, J. et al. Structure and receptor specificity of the hemagglutinin from an H5N1 influenza virus. Science 312, 404–410 (2006). 10.1126/science.1124513

8 Lin, T.-H. et al. A single mutation in bovine influenza H5N1 hemagglutinin switches specificity to human receptors. Science 386, 1128–1134 (2024). 10.1126/science.adt0180

9 Morano, N. C. et al. Structure of a zoonotic H5N1 hemagglutinin reveals a receptor-binding site occupied by an auto-glycan. Structure 33, 228–233.e223 (2025). 10.1016/j.str.2025.01.001

10 Ren, J. M., Rejtar, T., Li, L. & Karger, B. L. N-Glycan Structure Annotation of Glycopeptides Using a Linearized Glycan Structure Database (GlyDB). J. Proteome Res. 6, 3162–3173 (2007). 10.1021/pr070111y

11 Song, H. et al. Receptor binding, structure, and tissue tropism of cattle-infecting H5N1 avian influenza virus hemagglutinin. Cell 188, 919–929.e919 (2025). 10.1016/j.cell.2025.01.019

12 Gu, M. et al. The T160A hemagglutinin substitution affects not only receptor binding property but also transmissibility of H5N1 clade 2.3.4 avian influenza virus in guinea pigs. Vet. Res. 48, 7 (2017). 10.1186/s13567-017-0410-0

13 Pulit-Penaloza, J. A. et al. Transmission of a human isolate of clade 2.3.4.4b A(H5N1) virus in ferrets. Nature 636, 705–710 (2024). 10.1038/s41586-024-08246-7

14 Liu, M. et al. Human-type sialic acid receptors contribute to avian influenza A virus binding and entry by hetero-multivalent interactions. Nat. Commun. 13, 4054 (2022). 10.1038/s41467-022-31840-0

15 Kallewaard, N. L. et al. Structure and Function Analysis of an Antibody Recognizing All Influenza A Subtypes. Cell 166, 596–608 (2016). 10.1016/j.cell.2016.05.073

16 Guo, Y. et al. Analysis of hemagglutinin-mediated entry tropism of H5N1 avian influenza. Virol. J. 6, 39 (2009). 10.1186/1743-422x-6-39

17 Morse, J. et al. Influenza A(H5N1) Virus Infection in Two Dairy Farm Workers in Michigan. N. Engl. J. Med. 391, 963–964 (2024). 10.1056/NEJMc2407264

18 Maines, T. R. et al. Avian influenza (H5N1) viruses isolated from humans in Asia in 2004 exhibit increased virulence in mammals. J. Virol. 79, 11788–11800 (2005). 10.1128/jvi.79.18.11788-11800.2005

19 Oner, A. F. et al. H5N1 Avian Influenza in Children. Clin. Infect. Dis. 55, 26–32 (2012). 10.1093/cid/cis295

20 Swan, J. et al. The Sialome of the Retina, Alteration in Age-Related Macular Degeneration Pathology, and Potential Impacts on Complement Factor H. Invest. Ophthalmol. Vis. Sci. 66, 81 (2025). 10.1167/iovs.66.6.81

21 Herfst, S. et al. Airborne Transmission of Influenza A/H5N1 Virus Between Ferrets. Science 336, 1534–1541 (2012). 10.1126/science.1213362

22 Imai, M. et al. Experimental adaptation of an influenza H5 HA confers respiratory droplet transmission to a reassortant H5 HA/H1N1 virus in ferrets. Nature 486, 420–428 (2012). 10.1038/nature10831

23 Healy, C. M. Risk of Highly Pathogenic Avian Influenza A/H5N1 Virus in Pediatrics. J. Pediatr. Infect. Dis. Soc., piaf035 (2025). 10.1093/jpids/piaf035

24 Belachew, A. B., Rantala, A. K., Jaakkola, M. S., Hugg, T. T. & Jaakkola, J. J. K. Asthma and Respiratory Infections From Birth to Young Adulthood: The Espoo Cohort Study. Am. J. Epidemiol. 192, 408–419 (2022). 10.1093/aje/kwac210

25 Jha, A. et al. Patterns of systemic and local inflammation in patients with asthma hospitalised with influenza. Eur. Respir. J. 54, 1900949 (2019). 10.1183/13993003.00949-2019

26 Juhn, Y. J. Risks for infection in patients with asthma (or other atopic conditions): Is asthma more than a chronic airway disease? J. Allergy Clin. Immunol. 134, 247–257.e243 (2014). 10.1016/j.jaci.2014.04.024

27 Message, S. D. et al. Rhinovirus-induced lower respiratory illness is increased in asthma and related to virus load and Th1/2 cytokine and IL-10 production. Proc. Natl. Acad. Sci. U. S. A. 105, 13562–13567 (2008). 10.1073/pnas.0804181105

28 Andreev, K. et al. Antiviral Susceptibility of Highly Pathogenic Avian Influenza A(H5N1) Viruses Circulating Globally in 2022-2023. J. Infect. Dis. 229, 1830–1835 (2024). 10.1093/infdis/jiad418

29 Signore, A. V. et al. Neuraminidase reassortment and oseltamivir resistance in clade 2.3.4.4b A(H5N1) viruses circulating among Canadian poultry, 2024. Emerg. Microbes Infect. 14, 2469643 (2025). 10.1080/22221751.2025.2469643

30 McKimm-Breschkin, J. L. Influenza neuraminidase inhibitors: antiviral action and mechanisms of resistance. Influenza Other Respir. Viruses 7, 25–36 (2013). 10.1111/irv.12047

31 Li, Z. N. et al. Pre-existing cross-reactive immunity to highly pathogenic avian influenza 2.3.4.4b A(H5N1) virus in the United States. Nat. Commun. 16, 10954 (2025). 10.1038/s41467-025-66431-2

32 Arroyave, A. et al. Assessment of Cross-Reactive Neutralizing Antibodies Induction Against H5N1 Clade 2.3.4.4b by Prior Seasonal Influenza Immunization in Retail Workers. Open Forum Infect. Dis. 12, ofaf463 (2025). 10.1093/ofid/ofaf463

33 Ahmed, M. S. et al. Cross-reactive immunity against influenza viruses in children and adults following 2009 pandemic H1N1 infection. Antivir. Res. 114, 106–112 (2015). 10.1016/j.antiviral.2014.12.008

34 Lee, P. S., Zhu, X., Yu, W. & Wilson, I. A. Design and Structure of an Engineered Disulfide-Stabilized Influenza Virus Hemagglutinin Trimer. J. Virol. 89, 7417–7420 (2015). 10.1128/jvi.00808-15

35 Whittle, J. R. R. et al. Flow Cytometry Reveals that H5N1 Vaccination Elicits Cross-Reactive Stem-Directed Antibodies from Multiple Ig Heavy-Chain Lineages. J. Virol. 88, 4047–4057 (2014). 10.1128/jvi.03422-13

36 Pettersen, E. F. et al. UCSF ChimeraX: Structure visualization for researchers, educators, and developers. Protein Sci. 30, 70–82 (2021). 10.1002/pro.3943

37 Liebschner, D. et al. Macromolecular structure determination using X-rays, neutrons and electrons: recent developments in Phenix. Acta Crystallogr. D 75, 861–877 (2019). 10.1107/S2059798319011471

38 Emsley, P., Lohkamp, B., Scott, W. G. & Cowtan, K. Features and development of Coot. Acta Crystallogr. D D66, 486–501 (2010). 10.1107/S0907444910007493

39 Mehta, A. Y. & Cummings, R. D. GlycoGlyph: a glycan visualizing, drawing and naming application. Bioinformatics 36, 3613–3614 (2020). 10.1093/bioinformatics/btaa190

